# Raising animals without antibiotics: producer and veterinarian experiences and opinions

**DOI:** 10.1101/600965

**Authors:** Randall S. Singer, Leah J. Porter, Daniel U. Thomson, Mallory Gage, Amanda Beaudoin, Jennifer K. Wishnie

## Abstract

Ensuring the safety, health, and overall well-being of animals raised for food is both an ethical obligation and a critical component of providing safe food products. The use of antibiotics for maintaining animal health has come under scrutiny in recent years due to the rise of antibiotic resistance globally. Some U.S. producers, especially in the poultry industry, have responded by eliminating their antibiotic use. The number of animals raised without antibiotics (RWA) is growing in the U.S., but there are concerns that RWA practices might negatively impact animal health and welfare. Therefore, the objective of this study was to survey U.S. veterinarians and producers about their experiences and opinions regarding RWA production. Veterinarians, farmers, ranchers, producers, and other stakeholders involved in raising broilers, turkeys, swine, beef cattle or dairy cattle were surveyed. Of the 565 completed responses received, 442 self-reported as practicing veterinarians or producers. Just over half of respondents reported having past or current experience with RWA programs. The main indicated reasons for raising animals without antibiotics were market driven; switching to RWA production was less commonly made for health-related reasons, such as to reduce antibiotic resistance or to improve animal health and welfare. Although respondents felt that RWA production has negative impacts on animal health and welfare, they overwhelmingly (>70%) indicated that the customer (retailer/restaurant/food service) believes that animal and health welfare will be significantly improved. Veterinarians and producers indicated that RWA programs will increase production costs with questionable effect on meat, egg or dairy consumer demand. Many respondents felt that there are times when the RWA label takes priority over animal health and welfare. Respondents generally felt that there was a need for increased auditing/assessment of animal health and welfare in RWA systems.

## Introduction

Ensuring the health and well-being of animals raised for food is both an ethical obligation and a critical component of providing safe food products. Antibiotics are an important part of animal health programs, but their use has come under scrutiny because of the rise of antibiotic resistance globally (1-4). Efforts have been made to improve antibiotic stewardship in animal agriculture, with different countries often adopting different approaches for enhancing the responsible use of antibiotics (1, 5, 6).

Some animal producers, particularly within the U.S. poultry industry, have eliminated antibiotic use entirely and have adopted a “no antibiotics ever” (NAE) or “raised without antibiotics” (RWA) approach to animal production (7). In this paper we will refer to these programs as RWA. In RWA programs, antibiotics cannot be given to any of the animals via feed, water or by injection at any point in their lives; ill animals needing antibiotic therapy must be removed from the RWA program. These animals that received antibiotic therapy cannot be sold under an RWA label and must be marketed through a different distribution channel (8). Such circumstances often raise logistical challenges and potential financial losses for the producer.

RWA programs are intended to supply customers, such as restaurants, grocers and other food service establishments, with meat, eggs, and dairy products that can be labeled as having never had exposure to antibiotics. Anecdotal evidence suggests that retail customers and consumers assume that RWA and organic production will improve food safety and decrease antibiotic resistance in animals and humans while providing a more wholesome food product (9). In a recent survey of consumers, 55% responded that they were extremely or very concerned about antibiotic use in chickens when they purchase chicken (10). This same survey found that respondents generally had major misunderstandings about poultry production practices. For example, 60% of respondents considered themselves to be very or somewhat knowledgeable about the care of chickens, but 75% believed that there are added hormones or steroids in chicken meat (which has been illegal in the U.S. for many decades), and 71% believed that chickens raised for meat are housed in cages (which is untrue). Commercial broiler chickens are typically reared in enclosed housing that limits exposure to predators and disease-carrying wild birds and animals, and allows more control over environmental conditions. Birds range freely within the house. Over half of survey respondents disagreed with the statement “Eliminating antibiotics leads to significantly more chickens dying of disease.” While educating consumers is not the focus of this study, there is clearly a need to inform consumers about food animal production, which would necessitate transparency about production practices in the food animal sector and an openness by the consumer to receive this information from the food animal industry.

Few reports exist comparing RWA to conventionally-reared animals, particularly with respect to potential impacts on animal health, productivity, and welfare. A report was published in 2011 by Smith discussing his 12-year experience with RWA in broiler chickens (11), and some of his experiences included that these birds were more expensive to produce, due in part to stricter and more expensive diet requirements, and that the drug-free birds had a higher incidence of important diseases such as necrotic enteritis. More recently, Gaucher et al. (12) reported that drug-free production was associated with overall negative effects on key performance and gut health indicators (increased necrotic enteritis incidence, increased feed conversion, decreased daily weight gain, and decreased mean live slaughter weight), findings which are indicative of potentially negative impacts on overall animal welfare. These outcomes can contribute to economic and environmental strain, as RWA programs try to match production output of conventional programs.

A recent study compared three different broiler production systems: conventional, RWA, and non-medically important, wherein only antibiotics not considered important to human health are used (13). The study considered three important health conditions (eye ammonia burns, footpad lesions, and airsacculitis) which can be indicators of poor animal welfare. Pain from these conditions can lead to decreased feed intake and reduced weight gain. RWA production was shown to increase the risk and severity of all three of these health conditions. Use of non-medically important antibiotics diminished this risk and severity, but the risk was still higher and disease more severe than that in conventional systems. Study authors emphasized important limitations to their approach. First, the analyses do not prove a cause and effect relationship; in other words, the authors are not stating that raising birds RWA causes these conditions to become worse. Second, they emphasize that they did not analyze management practices and other related on-farm variables. They state that shifting to RWA production necessitates changes to production, such as reduced stocking density and longer downtime between flock production cycles in a barn. Thus many of the negative impacts of RWA production can potentially be diminished over time, but some might never be completely eliminated. For example, a recent randomized controlled trial in pigs found that animals reared under RWA conditions had worsened animal health when there were endemic viral and secondary bacterial infections on-farm (14).

As more animal production shifts from conventional to RWA programs, there is a need to understand the impacts of RWA systems on animal health and welfare. Because of the paucity of information regarding these potential impacts, we believe a survey of individuals directly involved with the raising of animals in the U.S. is needed to understand their experiences and opinions with RWA and conventional production. Therefore, the objective of this study was to survey veterinarians and producers directly involved in animal production about their experience and perception of the impacts (positive or negative) of RWA animal production on animal health and welfare. Specifically, this manuscript focuses on the effects of RWA production in the poultry, beef, swine, and dairy sectors on animal welfare, food safety, and cost of production.

## Materials and methods

### Survey design

The survey was designed to collect information from veterinarians and producers involved with beef cattle, dairy cattle, swine, turkey, and broiler chicken production. The survey tool was developed by study co-authors and was reviewed by industry experts in each commodity for clarity, completeness, and usability.

Respondents to the survey were only allowed to answer questions for one of the five animal commodities, and this was based on the commodity that the respondent selected at the very beginning of the survey as the commodity with which they were most familiar. The overall survey included questions related to the respondent’s RWA program experience, disease and welfare challenges within the respondent’s selected commodity, and experiences/beliefs about RWA impacts on animal health and welfare, food safety, cost of production, and antibiotic resistance. The survey was created for online administration using web-based survey software (Qualtrics, Provo, UT, USA) and collected no identifying information from respondents. A complete print-version of the survey is included in S1 Appendix.

### Survey dissemination

A hyperlink to the online survey was distributed by various professional organizations and commodity groups such as American Association of Avian Pathologists (AAAP), National Chicken Council (NCC), National Turkey Federation (NTF), U.S. Poultry & Egg Association (USPOULTRY), American Association of Bovine Practitioners (AABP), Academy of Veterinary Consultants (AVC), Animal Agriculture Alliance, National Pork Producers Council (NPPC), National Pork Board (NPB), American Association of Swine Veterinarians (AASV), and Pig Improvement Company (PIC). Announcements were also made at multiple professional and commodity meetings and in key trade journals. The survey was open from February 15 to March 23, 2018.

### Data analysis

Incomplete surveys were excluded from analysis. This survey was intended to focus on animal production within the U.S. Because of the potential for varying regulation, management practices and production systems to influence responses, data from international respondents were excluded from analysis. Data analysis was conducted using standard statistical software (Stata 15.1, College Station, TX, USA). Respondents were categorized as having any experience with RWA production (RWA respondent) or having no experience with RWA production (Conventional respondent). Respondent role (e.g., veterinarian, producer) and RWA experience were compared with two-sample Wilcoxon rank-sum (Mann-Whitney) tests. Likert scale graphs were prepared in R (15) using packages licorice and ggplot2 (16).

Analyses in this paper focus on study questions related to potential impacts of RWA production on food safety, animal welfare, cost of production, demand for the respondent’s animal protein or product, and auditing of RWA production systems. Study questions that focused on impacts on specific animal diseases, animal production, and disease interventions are not addressed.

## Results

### Survey responses

Five hundred and sixty-five completed responses were received, although not every question was necessarily answered. Ninety-five percent of respondents (n=536) were located within the U.S. (Table 1). Twenty-seven international respondents were excluded from the analysis and are not included in the results that follow. Most respondents were practicing veterinarians (n=248, 43.9%), producers (n=214, 37.9%), and technical services professionals (n=44, 7.8%). Just over half of the respondents were working with (n=241, 42.7%) or had previously worked with (n=76, 13.5%) animals being raised without antibiotics (RWA respondents). The remaining respondents (n=248, 43.9%) had no direct experience with RWA production (Conventional respondents). For the following analyses, only producers and veterinarians with direct animal responsibilities are included (i.e. technical services professionals, academics and government employees are excluded). Because only one turkey respondent had no experience with RWA production, no details of this response are provided. A total of 442 responses are included in the analyses that follow, although no information is provided about the single Conventional turkey respondent. These 442 respondents had completed all or most of the questions addressed in this manuscript

**Table 1:** Characteristics of survey respondents, n=565.

|  | **Total** | **Broiler** | **Turkey** | **Swine** | **Beef** | **Dairy** |
| --- | --- | --- | --- | --- | --- | --- |
| **Role** | **565** | **69** | **23** | **148** | **244** | **81** |
| Practicing Veterinarian | 43.9% | 31.9% | 52.2% | 37.6% | 43.4% | 64.2% |
| Research/Academic/Government Veterinarian | 5.1% | 1.5% | 4.4% | 4.7% | 4.1% | 12.4% |
| Research/Academic/Government Non-veterinarian | 1.1% | 2.9% | - | 0.7% | 1.2% | - |
| Manager/Producer/Grower/Rancher/Owner | 37.9% | 26.1% | 26.1% | 47.3% | 44.3% | 14.8% |
| Technical Services | 7.8% | 29.0% | 13.0% | 5.4% | 2.9% | 7.4% |
| Other | 4.3% | 8.7% | 4.4% | 4.1% | 4.1% | 1.2% |
| **Country of Experience** |  |  |  |  |  |  |
| United States | 95.2% | 86.8% | 95.8% | 96.0% | 97.5% | 92.6% |
| International | 4.8% | 13.2% | 4.2% | 4.1% | 2.5% | 7.4% |
| **Experience with RWA** |  |  |  |  |  |  |
| Current Experience | 42.7% | 63.8% | 95.7% | 33.8% | 36.1% | 45.7% |
| Previous Experience | 13.5% | 2.9% | - | 20.3% | 13.5% | 13.6% |
| No Experience | 43.9% | 33.3% | 4.4% | 46.0% | 50.4% | 40.7% |

Respondents indicated the factors that contributed to their decision to participate in RWA production (RWA respondents) or reasons why they did not (Conventional respondents), and these responses are shown in Table 2. RWA respondents in all commodities most commonly identified market-driven reasons for their decision to participate in RWA production. Specifically, the most common reason was “to fulfill a client/customer request” (>60% across all commodities). Conventional respondents most commonly identified “concerns about negative impacts to animal health and welfare” (>60% across all commodities) and “already raising animals in a responsible [antibiotic] use program” (>50% across all commodities) as the most common reasons for not participating in RWA production.

**Table 2:** Factors contributing to decision to raise animals RWA or conventionally, n=442.

|  | **Broiler** | **Turkey** | **Swine** | **Beef** | **Dairy** |
| --- | --- | --- | --- | --- | --- |
| **RWA Respondents** | **19** | **17** | **59** | **97** | **36** |
| To decrease antibiotic resistance | 26.3% | 5.9% | 8.5% | 21.6% | 2.8% |
| To improve animal health and welfare | 26.3% | 5.9% | 10.2% | 17.5% | 8.3% |
| To increase sale price of animals/product | 42.1% | 41.2% | 62.7% | 41.2% | 11.1% |
| To gain market entry into a retail program | 36.8% | 58.8% | 37.3% | 27.8% | 8.3% |
| To fulfill a client/customer request | 84.2% | 82.4% | 69.5% | 62.9% | 77.8% |
| To eliminate the use of medically important antibiotics | 10.5% | 0.0% | 5.1% | 11.3% | 5.6% |
| **Conventional Respondents** | **15** | **1** | **63** | **110** | **25** |
| Not profitable | 33.3% | - | 27.0% | 22.7% | 8.0% |
| Concerned about negative impacts to animal health and welfare | 93.3% | - | 76.2% | 68.2% | 68.0% |
| No market pressure | 20.0% | - | 31.7% | 26.4% | 24.0% |
| Not a sustainable consumer trend | 40.0% | - | 25.4% | 13.6% | 8.0% |
| Food safety concerns | 13.3% | - | 30.2% | 8.2% | 24.0% |
| Already raising animals in a responsible use program | 60.0% | - | 71.4% | 57.3% | 68.0% |

### Animal health and welfare

Respondents were asked how they thought RWA production impacts animal health and welfare. Across all five commodities, most RWA and Conventional respondents (> 60% for all commodities) believed that RWA production would slightly worsen or significantly worsen animal health and welfare (Fig 1). Within the broiler, beef, and swine responses, significantly more Conventional respondents believed that RWA production would negatively impact animal welfare than did RWA respondents (P<0.01, P<0.01, and P<0.05, respectively); there was no statistically significant difference between Conventional and RWA dairy respondents. Among RWA respondents, producers perceived less of a negative impact on animal health and welfare than did veterinarians. Conventional veterinarian and producer perceptions were more aligned, with both believing that the animal health and welfare impact would be more negative than the beliefs of their RWA counterparts.

**Fig 1:**
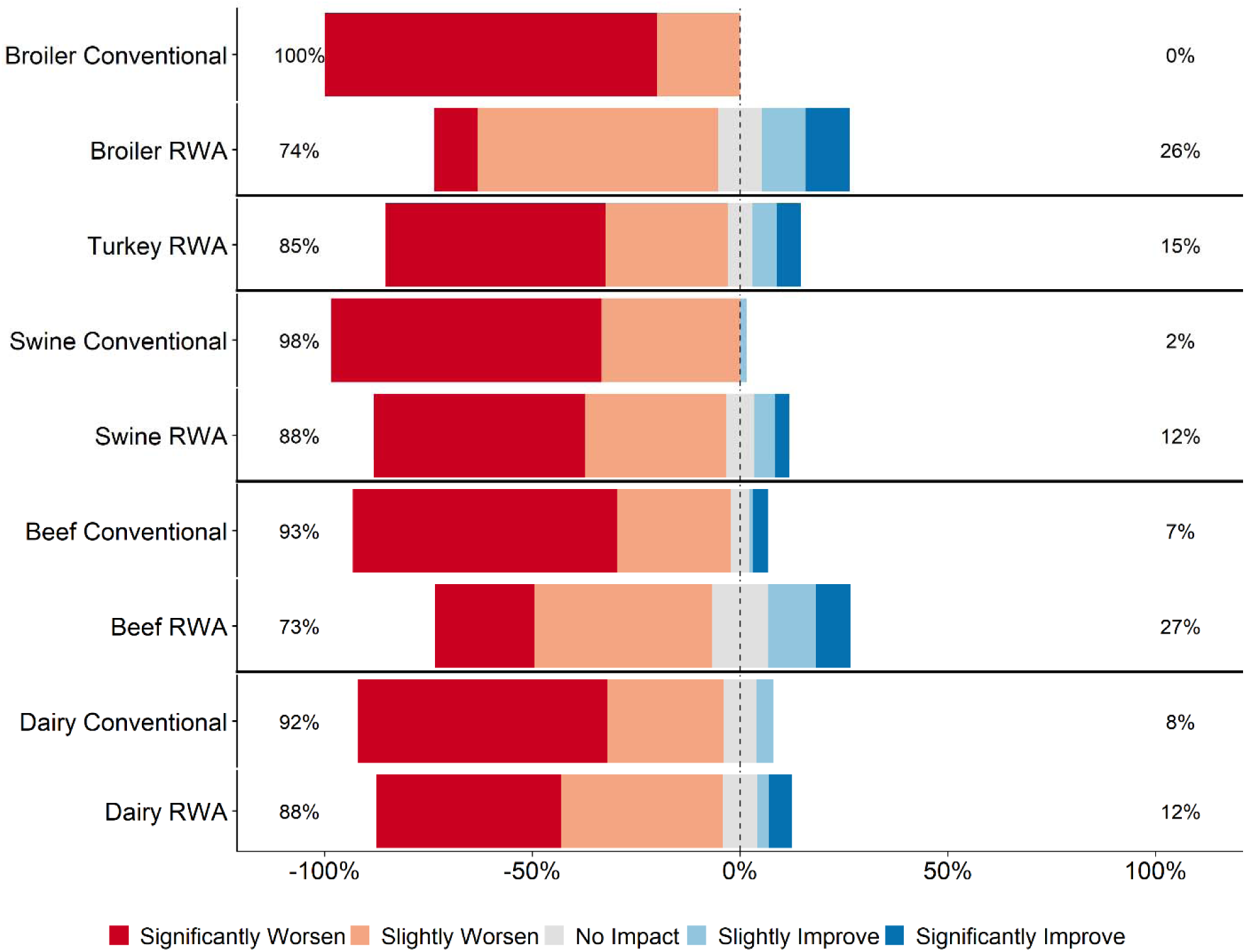
Respondents’ opinion about impact of RWA production on animal health and welfare. Five-item Likert scale reporting respondents’ opinion, stratified by commodity and RWA experience.

Respondents were asked for their perception of customer (retailers, restaurants, or food services) opinions regarding how RWA production impacts animal health and welfare. The perception of the majority of RWA and Conventional respondents (> 60% for all commodities) was that their customers believe that raising animals without antibiotics would slightly improve or significantly improve animal health and welfare (Fig 2). This perception did not differ between RWA and Conventional respondents.

**Fig 2:**
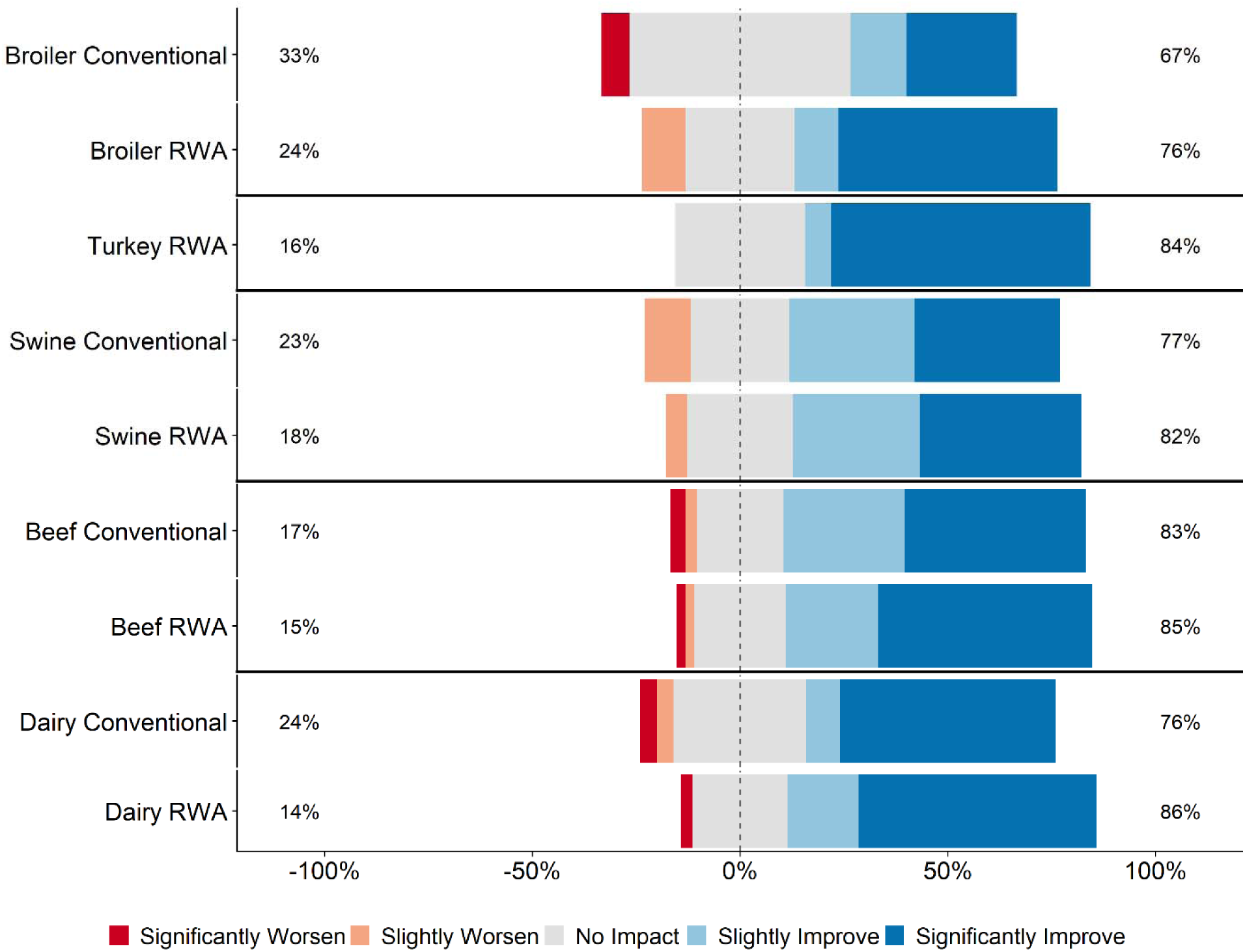
Respondents’ opinion about customer perception regarding the impact of RWA production on animal health and welfare. Five-item Likert scale reporting respondents’ opinion, stratified by commodity and RWA experience.

### Food safety

Across all five commodities, the majority of RWA and Conventional respondents (> 55% for all commodities except RWA beef respondents at 45%) believed that raising animals without antibiotics would have no impact, slightly worsen or significantly worsen food safety (Fig 3). Within the broiler and beef responses, significantly more Conventional respondents believed that RWA production would negatively impact food safety than did RWA respondents (P<0.01 for broiler and beef). When stratified by role, there was a difference of opinion in the RWA respondent group between veterinarians and producers, with RWA producers believing that there would be less of a negative impact on food safety when antibiotics are removed from the production system than did RWA veterinarians. Within the Conventional group of respondents, veterinarian and producer perceptions were more aligned regarding the impact of removing antibiotics from the production system on food safety.

**Fig 3:**
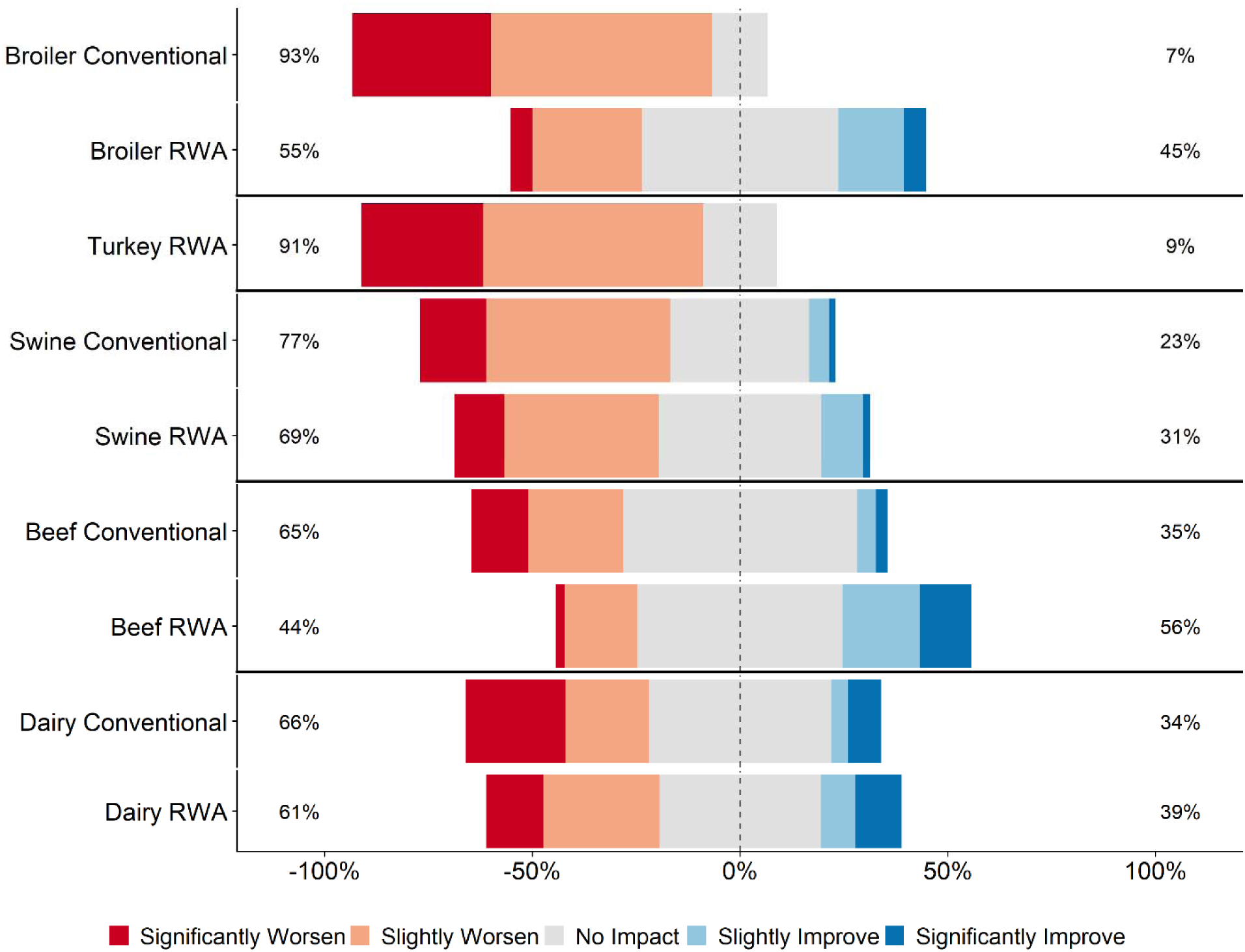
Respondents’ opinion about the impact of RWA production on food safety. Five-item Likert scale reporting respondents’ opinion, stratified by commodity and RWA experience.

Across all five commodities, the perception among the majority of RWA and Conventional respondents (> 60% for all commodities) was that their customers (retailers, restaurants, or food services) believed that raising animals without antibiotics would slightly improve or significantly improve food safety (Fig 4). There were no statistically significant differences between RWA and Conventional veterinarians or producers within any of the commodities; there was a general perception that customers believe that food safety is improved by RWA production practices.

**Fig 4:**
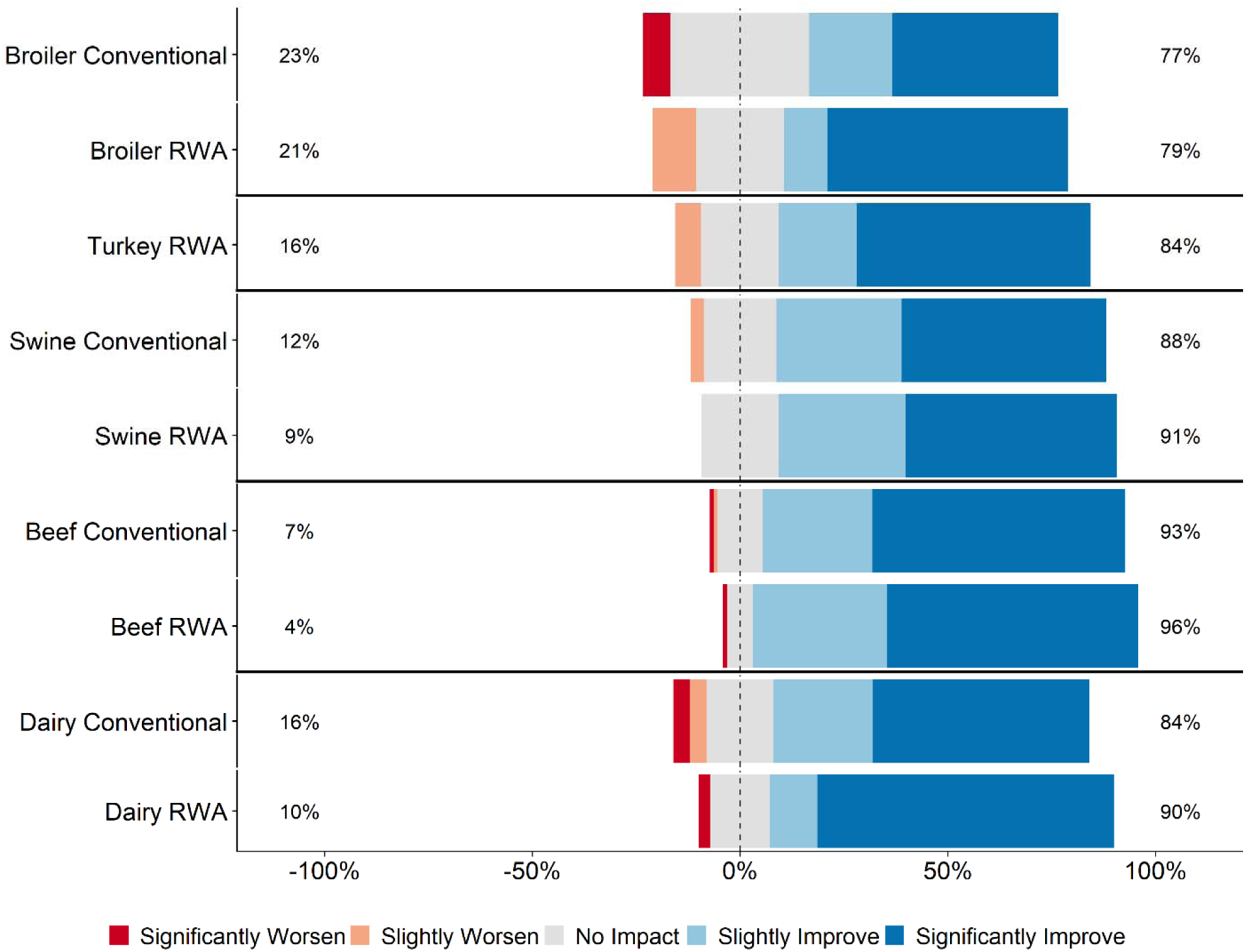
Respondents’ opinion about customer perception regarding the impact of RWA production on food safety. Five-item Likert scale reporting respondents’ opinion, stratified by commodity and RWA experience.

### Cost and demand

Across all five commodities, most RWA and Conventional respondents (> 80%) believed that raising animals without antibiotics would slightly or significantly increase the cost of production (Fig 5). Among those respondents that work with beef cattle, significantly more Conventional respondents believed that the cost of production would be increased than did RWA respondents (P<0.01); there were no statistically significant differences within the other commodities. Across all five commodities and RWA experiences, veterinarians were more likely than producers to say that production costs would be increased.

**Fig 5:**
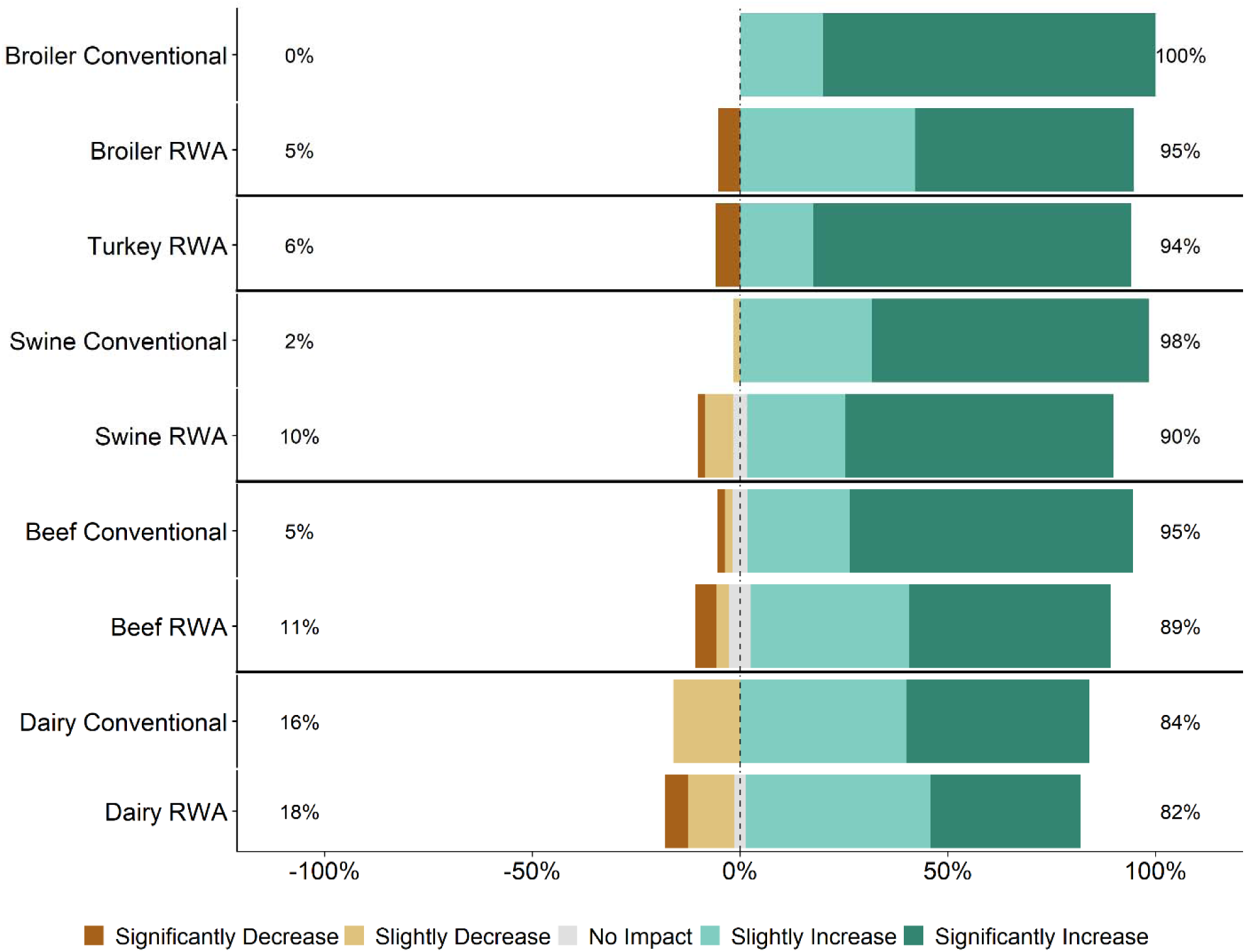
Respondents’ opinion about the impact of RWA production on cost of production. Five-item Likert scale reporting respondents’ opinion, stratified by commodity and RWA experience.

Respondents were also asked how they think RWA production would impact demand for their protein or product. Across all five commodities, most RWA and Conventional respondents (> 80%) believed that raising animals without antibiotics would have no impact or would slightly increase demand for their protein (Fig 6). Significantly more beef, dairy, and broiler RWA respondents believed that demand would be increased when compared to Conventional respondents (P<0.05 for each commodity). Across all five commodities and RWA experiences, producers were more likely than veterinarians to say that the demand for the protein or product would be increased.

**Fig 6:**
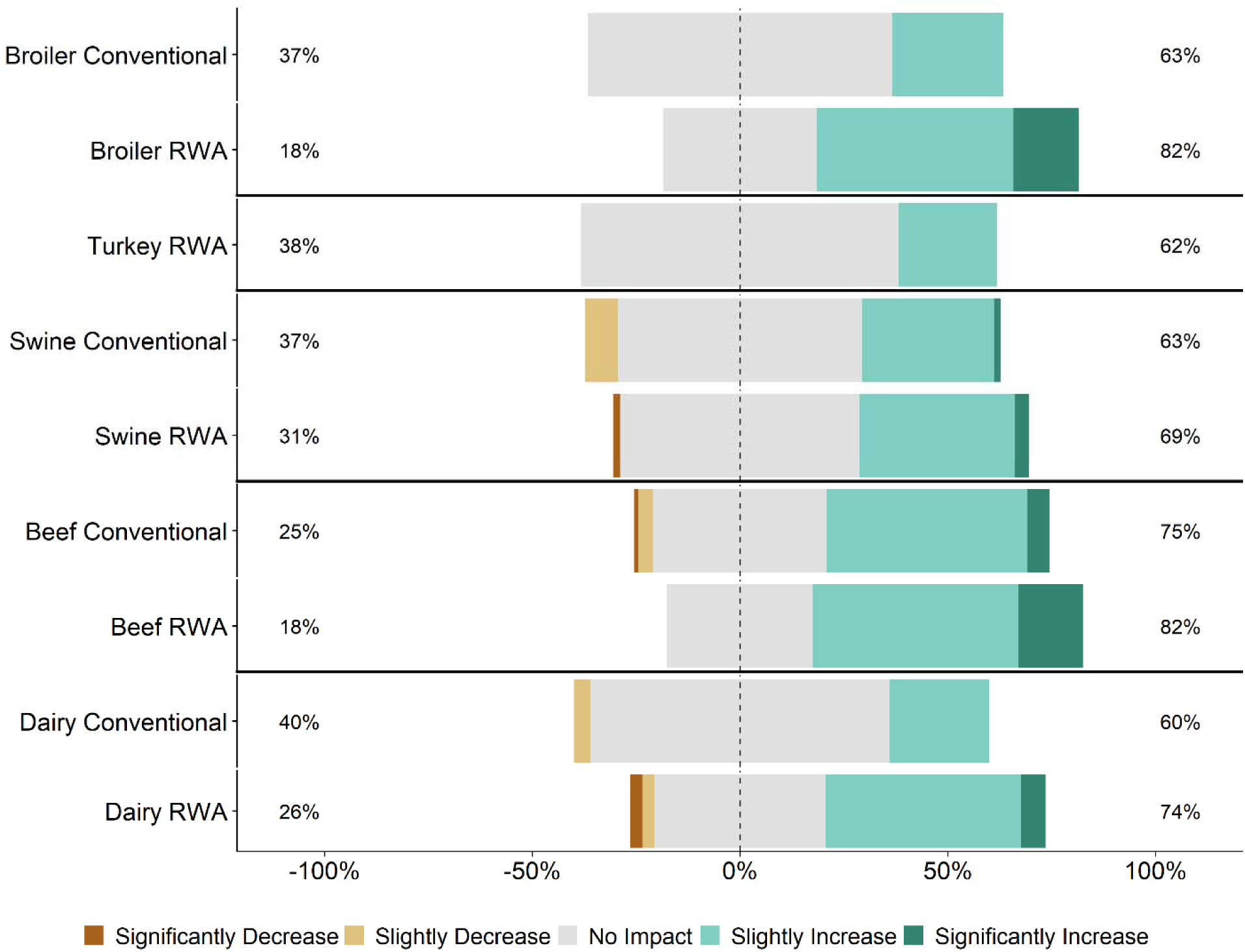
Respondents’ opinion about the impact of RWA production on demand for their commodity’s protein or product. Five-item Likert scale reporting respondents’ opinion, stratified by commodity and RWA experience.

### Label and auditing

Respondents were asked whether maintaining the RWA label on a product ever takes priority over flock/herd health and welfare. Specifically, survey participants were asked how strongly they agree or disagree with the statement: “There are times that maintaining an RWA label has priority over flock/herd health and welfare.” Regardless of commodity type and RWA experience, responses to this question ranged from Strongly Disagree to Strongly Agree (Fig 7). A higher percentage of RWA swine and dairy respondents Somewhat Agreed or Strongly Agreed with this statement than Conventional respondents, whereas the percentages were approximately equal for the beef and broiler chicken respondents. In general, there were no major differences between the RWA and Conventional respondents when stratified by role. The analysis was repeated for the veterinarian respondents because the decision to use an antibiotic is made by the veterinarian and thus the veterinarian respondents should have a better ability to address this question of the survey. Regardless of commodity type and RWA experience, the veterinarian respondents again had a range of responses, including many who Somewhat Agreed or Strongly Agreed with the statement (Fig 8).

**Fig 7:**
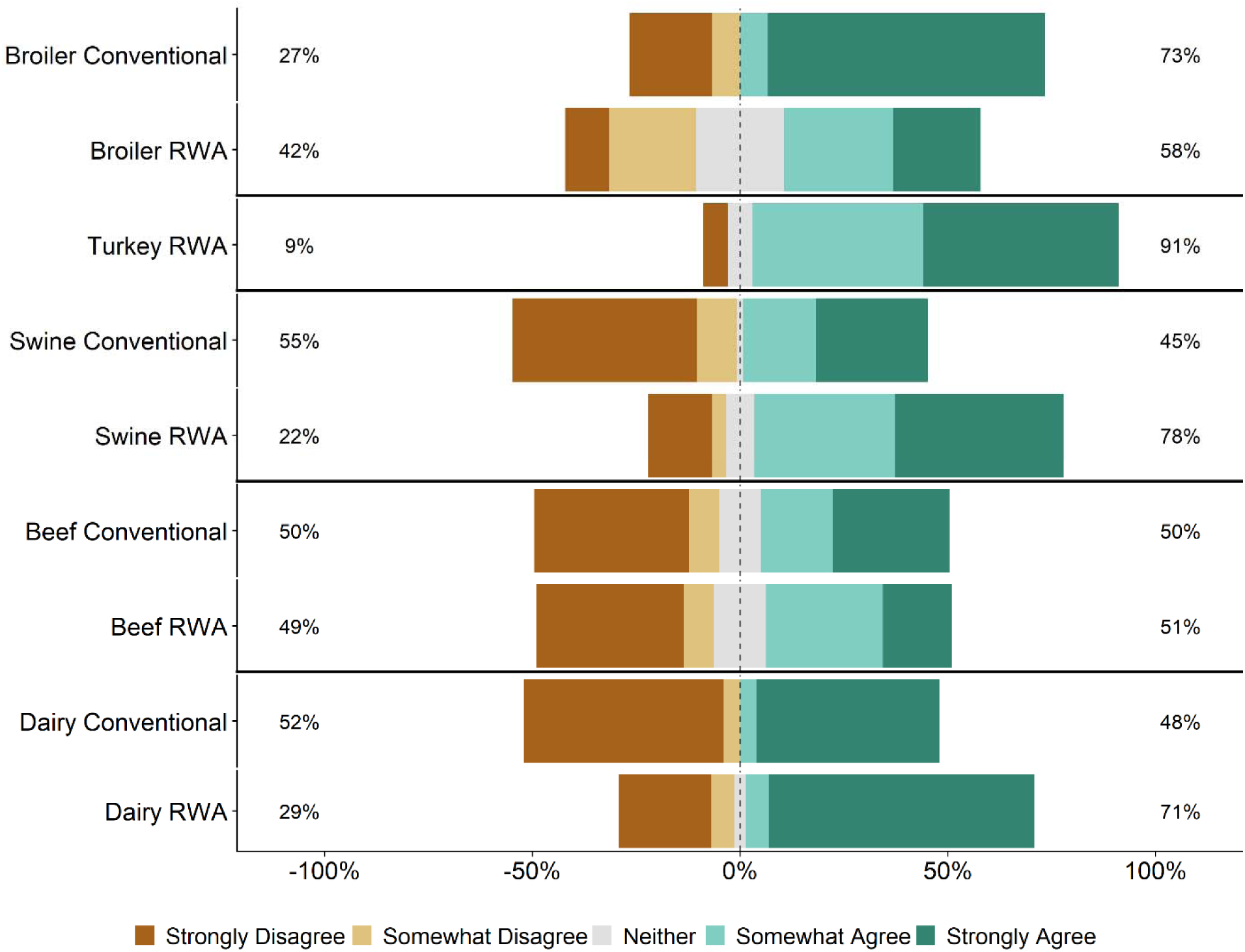
Respondents’ opinion about the statement, “There are times that maintaining a raised without antibiotics label has priority over flock/herd health and welfare.” Five-item Likert scale reporting respondents’ opinion, stratified by commodity and RWA experience.

**Fig 8:**
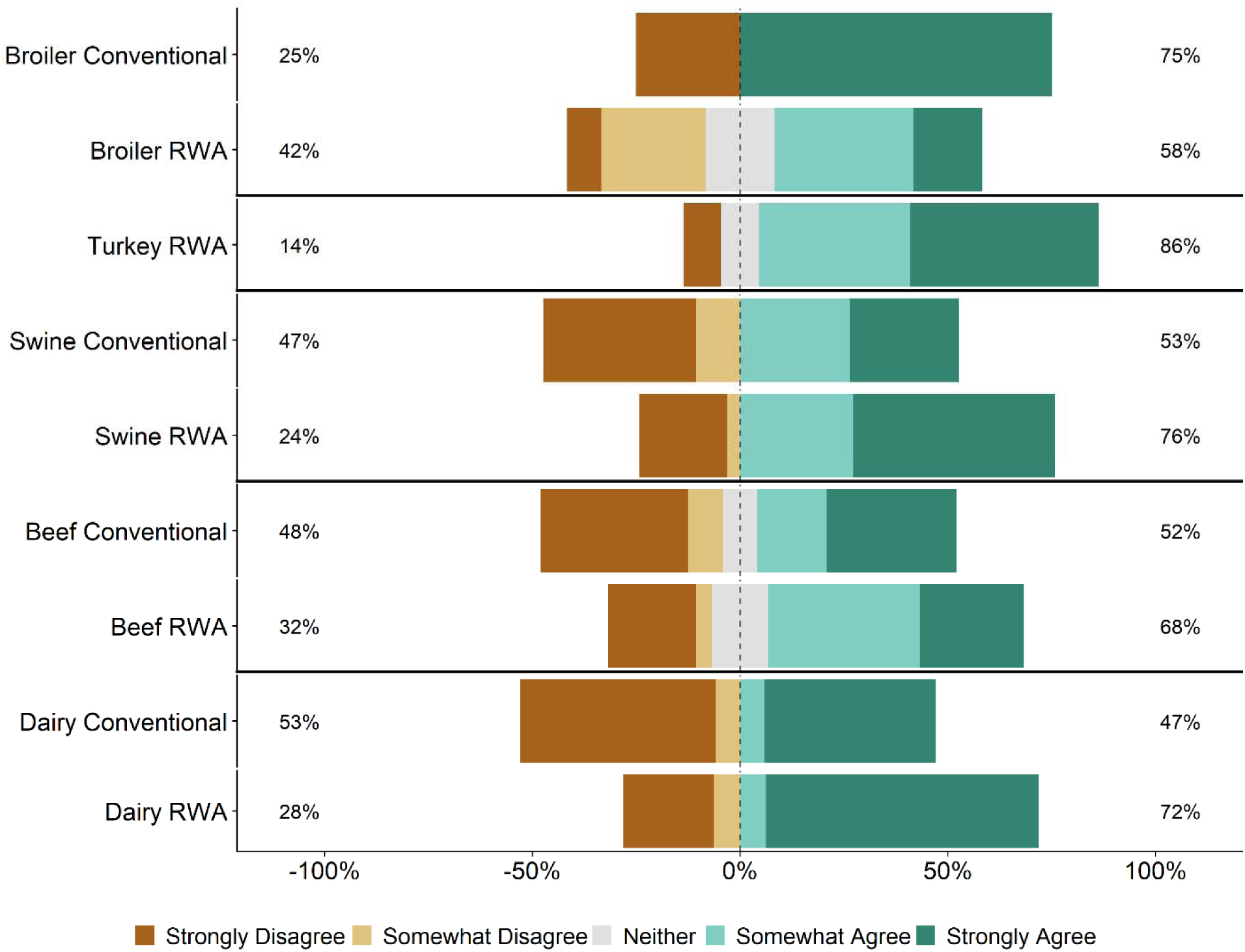
Veterinarian respondents’ opinion about the statement, “There are times that maintaining a raised without antibiotics label has priority over flock/herd health and welfare.” Five-item Likert scale reporting respondents’ opinion, stratified by commodity and RWA experience.

Respondents were asked whether more stringent health and welfare auditing and assessment is needed when raising animals without antibiotics. Across all five commodities and for both Conventional and RWA respondents, most respondents said that they Somewhat Agree or Strongly Agree with the need for more auditing and assessment in RWA settings with the exception of the RWA broiler respondents; only 32% of RWA Broiler respondents said that they Somewhat or Strongly Agree with this need (Fig 9). When stratified by role, Conventional veterinarians and producers were more likely to agree with the statement than the RWA veterinarians and producers.

**Fig 9:**
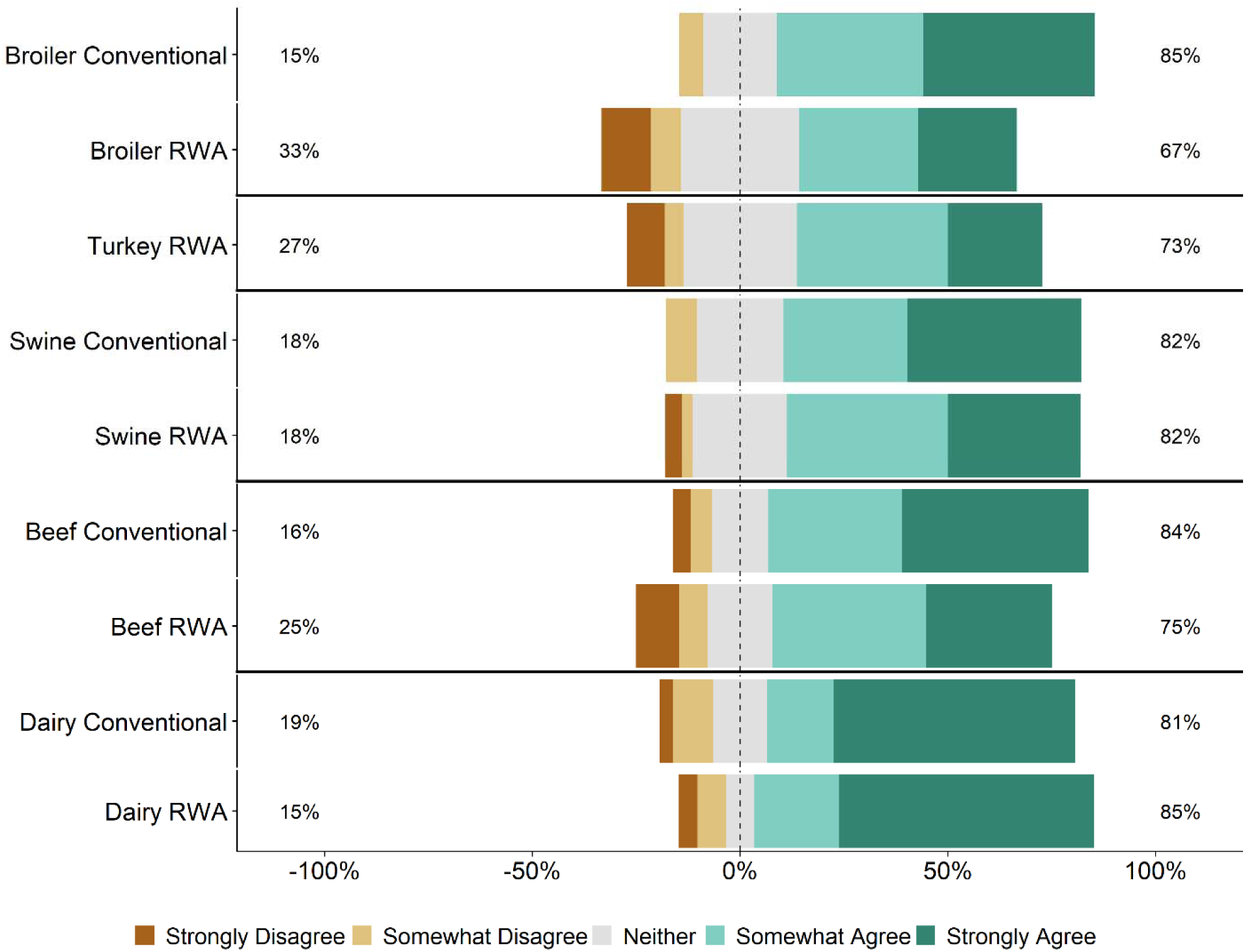
Respondents’ opinion about the need for more stringent health and welfare auditing/assessment when animals are raised without antibiotics. Five-item Likert scale reporting respondents’ opinion, stratified by commodity and RWA experience.

## Discussion

This survey was designed to gauge veterinarian and producer experiences and opinions regarding the impacts of RWA animal production on animal health and welfare. The main reasons for raising animals without antibiotics were market driven (Table 2), and in most circumstances, the decision to switch to RWA production was not made for health-improvement reasons, such as to reduce antibiotic resistance or to improve animal health and welfare. On the contrary, the RWA respondents generally tended to indicate that raising animals without antibiotics negatively affected animal health (Table 2 and Fig 1).

Veterinarians and producers indicated that RWA programs increase production costs (Fig 5) but were less certain that there would be a concomitant increase in consumer demand (Fig 6). Although respondents largely felt that RWA production negatively impacts animal health and welfare, they overwhelmingly share the perception that the customer (retailers, restaurants or food services) believes that animal health and welfare will be significantly improved by raising animals without antibiotics (Fig 2). Many respondents felt that there are times when maintaining the RWA label takes priority over animal health and welfare (Figs 7 and 8). In general, across all surveyed commodities, respondents saw a need for increased auditing and assessment of animal health and welfare in RWA systems (Fig 9).

Antibiotics remain an important component of health management in animal agriculture. The decision to use an antibiotic, including the optimization of when, why and for how long to administer the antibiotic, can be a complex and multi-faceted topic (17, 18). As is true in the varied settings and situations of human healthcare, approaches to improving antibiotic stewardship in animal agriculture, while effectively maintaining animal health and welfare, will differ among commodity types, animal operations and their veterinarians. A better understanding of the risks and benefits associated with RWA production is needed, in addition to the documentation of the changes that have been made in RWA systems to successfully maintain animal health and welfare. This current study helps fill some of these knowledge gaps and highlights areas where more information is needed.

This study has several key limitations. First, the study utilized an anonymous survey approach. As is the case with most surveys, particularly those that maintain the anonymity of respondents, it is impossible to follow up with the respondents to verify their responses. Regardless, this study was intended to fill a key data gap, namely the lack of data about RWA versus conventional production practices. The survey was used as a first step to gauge the perception of RWA animal production by those actively raising and those choosing not to raise animals in this system. A second possible limitation of this survey approach is the potential incentive of conventional respondents to overstate the negative aspects of RWA production. However, when viewed side-by-side for each commodity, the responses of the RWA and conventional participants are fairly consistent. Even though the RWA responses were based on the participants’ experience of RWA production, it would appear that the RWA and conventional respondents had similar perceptions of RWA production. Third, the questions regarding food safety may have been based on pure opinion for many of the respondents because it is unclear how much food safety information the producers and veterinarians receive about the animals under their care. Some producers and veterinarians receive feedback from the processing plant about the foodborne pathogen status of their animals, for example with respect to *Salmonella*, so while the respondents might not be experts on all food safety issues, they should have some information about certain foodborne pathogens in their animals.

The findings from this study indicate that the retailers, restaurants and food services might have a skewed perception of the impacts of RWA production. This is highlighted by the respondents’ opinions that their customers believe that RWA production improves animal health and welfare (Fig 2), in contrast to their own experiences and opinions (Fig 1). Studies of food industry customers are needed to determine the basis for their perceptions of the RWA impact on animal health and welfare and to better understand the systems used to audit RWA production. Importantly, a detailed assessment of the auditing that the customers do to ensure that animal health and welfare are being maintained in RWA systems is critical (19). If audits are conducted infrequently, on a small number of premises, or rely exclusively on the opinions and reports of the producers, it is possible that health and welfare problems would be missed. Clearly there is a need to educate customers and consumers about the role of antibiotics in food animal production and the challenges of eliminating antibiotics completely from the production system. Findings from this study can hopefully be used to advance this conversation.

The impacts of raising animals without antibiotics are not restricted to animal health and welfare. There are also potential effects on environmental sustainability and economic viability. One recent study developed a simulation model to evaluate the impacts of RWA broiler production (20). They estimated that if the entire U.S. broiler industry were to shift to RWA production, impacts would include decreased edible meat, an increase in the number of broilers needed to meet current demand (680-880 million more birds), associated increases in feed and water requirements (5.4-7.6 million excess tons and 1.9-3 billion excess gallons, respectively), and increased manure production (4.6-6.1 million excess tons). The authors conclude that “eliminating the use of antibiotics in the raising of broilers may have a negative effect on the conservation of natural resources as well as a negative economic effect via increased prices to the consumer. Results suggest the need to communicate to consumers the supportive role that prudent, responsible use of antibiotics for animal disease treatment, control, and prevention plays in the sustainable production of broilers.”

Animal health and welfare, and environmental and economic sustainability, are key considerations when evaluating RWA production. However, the initial motivation of RWA production was the goal of reducing antibiotic resistance of human and animal health importance. Analyses comparing resistant bacteria and resistance gene loads on conventional and RWA farms or mathematical modeling studies have reported conflicting results (21–23). There is a need for well-designed, longitudinal studies on farms that can simultaneously collect data on antibiotic use and resistance so that efforts to improve antibiotic stewardship can take resistance outcomes into account. It is important to have an evidence-based understanding of whether RWA production accomplishes the outcome for which it was intended, reducing antibiotic resistance on the farm.

Based on the responses to this survey, RWA production does not appear to be driven by prioritization of animal health and welfare. Many respondents felt that there are times when the RWA label takes priority over animal health and welfare. This observation is deeply concerning, as protecting animal health and welfare is a key component of the veterinarian’s oath (24). If animals receive antibiotics to treat disease, the meat from these animals cannot be marketed RWA, and the producers must absorb the added costs associated with RWA production. This might lead to pressures to sacrifice animal health and welfare to stay in an RWA program. As stated by Karavolias et al. (13), “Policies aimed at eliminating or restricting the use of antibiotics in broiler production may come with potentially negative consequences with respect to good animal welfare. A more effective policy approach should consider comprehensive animal care plans that incorporate good housing, management, and responsible antibiotic use, including the use of ionophores. Policies aimed at informing the consumer on the positive role of access to antibiotics in supporting good animal welfare while limiting risk of antibiotic resistance in humans are needed to address the current information gap.” Clearly educational programs are needed for the customers and consumers of meat products regarding animal production practices and the interventions that are available for maintaining animal health and welfare. Furthermore, veterinarians in animal agriculture need to continue to develop antibiotic stewardship programs to ensure that antibiotic use practices are optimized and that antibiotics are only being used when necessary.

## Acknowledgements

The authors thank Haejin Hwang for assistance with the R code used in the data analysis and Likert scale graphing.

## Author Contributions Statement

RS, DT, and JW conceived the study. RS, LP, DT, MG, and AB JW designed the survey. RS, LP, and JW analyzed the data. RS and LP prepared the initial draft, figures, tables, and appendices. All authors contributed to the writing and editing of the manuscript.

