## Supplementary material for "Raising animals without antibiotics: producer and veterinarian experiences and opinions": S1 Appendix

### NAE Welfare Survey - Final

#### Introduction

**Welcome to the Raised Without Antibiotics Survey**
Antibiotics are important for maintaining animal health, but their use has come under scrutiny in recent years due to the rise of antibiotic resistance globally. In the U.S., changes have been made to improve antibiotic stewardship in animal agriculture; some producers, especially in poultry, have responded by eliminating their antibiotic use altogether. Demand for poultry and livestock raised without the use of antibiotics is growing in the U.S., but there are few data available regarding the association between raised without antibiotic claims and animal health and welfare.

Our team has developed a survey to assess opinions and experiences of raised without antibiotics programs in animal agriculture and their relationship to animal health and welfare. This survey is intended to provide a detailed, evidence-based report that addresses the potential impacts that may arise from poultry and livestock raised without the use of antibiotics and will help to better inform consumers, producers, and other stakeholders about raised without antibiotics programs. **The survey is completely anonymous and confidential**; no data about individual participants are being recorded. Please be as honest as possible in your responses to the survey. Even if you have never raised animals without any antibiotics, your opinions are still needed.

This survey should take about 15 minutes to complete, and to repeat, **your participation is completely anonymous**. Survey results will only be reported in the aggregate thereby keeping your individual responses confidential. We appreciate your willingness to participate.

Sincerely,

Randall Singer (Mindwalk Consulting Group, LLC and University of Minnesota)
Dan Thomson (Thomson Livestock Consulting, LLC and Kansas State University)
Jennifer Wishnie (Wishnie Consulting, LLC and California Polytechnic University)
Mallory Gage (Gage Group Consulting, LLC)
Leah Porter (Mindwalk Consulting Group, LLC)
Amanda Beaudoin (Mindwalk Consulting Group, LLC and University of Minnesota)

Funding for this project is being provided by the Animal Agriculture Alliance. The project is being conducted independently by the investigators, who attest that the opinions and work contained herein accurately reflect their opinions and not necessarily those of the Animal Agriculture Alliance.

Q1.2 For which of the following commodities are you responding?
*(Please select the one for which you have the most experience)*

- Broiler (1)
- Turkey (2)
- Swine (3)
- Beef (4)
- Dairy (5)

---

Q1.3 What is your current role when working with this commodity?
(If you have more than one role please select the one you consider to be your primary role.)

- Practicing veterinarian (1)
- Research/Academic/Government veterinarian (2)
- Researcher/Academic/Government non-veterinarian (3)
- Manager/Producer/Grower/Rancher/Owner (4)
- Technical services (5)
- Other (please list) (6) ____________________

---

Q1.4 In what country do/did you work with this commodity?
*(If you have experience both in the U.S. and internationally, please consider the location with which you are most familiar for this survey.)*

- United States (1)
- Internationally (2)

### Broiler

Q2.2 Have you ever, past or present, produced/consulted/worked with broilers enrolled in marketing programs where the animals were raised without the use of any antibiotics?
- o Yes, I am currently working with broilers being raised without antibiotics
- o Yes, I have previously worked with broilers being raised without antibiotics but no longer do so
- o No, I have never worked with broilers raised without antibiotics

*Display This Question if NAE = YES PREVIOUSLY*

Q2.3 Why did you stop working with broilers raised without the use of any antibiotics?

*Display This Question if NAE = YES CURRENTLY or YES PREVIOUSLY*

Q2.4 Which of these factors contributed to your decision to produce/consult/work with broilers where the use of antibiotics was not allowed? (Select all that apply)
- o To decrease antibiotic resistance (1)
- o To improve animal health and welfare (2)
- o To increase sale price of animals/product (3)
- o To gain market entry into a retail program (4)
- o To fulfill a client/customer request (5)
- o To eliminate the use of medically important antibiotics (6)
- o Other (please list) (7) ________________________________________________

*Display This Question if NAE = NO*

Q2.5 Which of these factors contributed to your decision not to produce/consult/work with broilers where the use of antibiotics was not allowed?
(Select all that apply)
- o Not profitable (1)
- o Concerned about negative impacts to animal health and welfare (2)
- o No market pressure (3)
- o Not a sustainable consumer trend (4)
- o Food safety concerns (5)
- o Already eliminated the use of medically important antibiotics (6)
- o Already raising animals in a responsible use program (7)
- o Other (please list) (8) ________________________________________________

*Display This Question if NAE = YES CURRENTLY or YES PREVIOUSLY*

Q2.6 Is your experience in producing/consulting/working with broilers that have been raised without the use of antibiotics part of any of the following types of programs? (Select all that apply)
- o Industry sponsored program (1)
- o Private/Retail/Restaurant/Food Service program (2)
- o Packer/Processor program (3)
- o State/Federal government program (4)
- o Other (please list) (5) ________________________________________________
- o No program (6)

*Display This Question if NAE = NO*

Q2.7 Have you ever considered producing/consulting/working with broilers raised without the use of antibiotics as part of any of the following types of programs? (Select all that apply)

- Industry sponsored program (1)
- Private/Retail/Restaurant/Food Service program (2)
- Packer/Processor program (3)
- State/Federal government program (4)
- Other (please list) (5) ____________________
- No program (6)

---

Q2.8 Are the broilers you produce/consult/work with currently part of an animal welfare program? (Select all that apply)

Industry sponsored quality assurance program (1)

- Private/Retail/Restaurant/Food Service animal welfare program (2)
- Packer/Processor animal welfare program (3)
- State/Federal government animal welfare program (4)
- Other (please list) (5) ____________________
- No program (6)

---

*Display This Question if NAE = YES CURRENTLY or YES PREVIOUSLY*

Q2.9 Rank these disease challenges from <u>most to least</u> problematic when raising broilers without antibiotics. **(Click and drag to rank.)**

______ Necrotic enteritis (126)
______ Airsacculitis (E. *coli* respiratory infection) (127)
______ Gangrenous dermatitis (128)
______ Kinky back (spondylolisthesis) (129)
______ Cellulitis / IP (inflammatory process) (130)
______ Viral enteric disease (131)
______ Bacterial osteomyelitis / lameness (132)
______ Other *(please list)* (133)

---

*Display This Question if NAE = NO*

Q2.10 Rank these disease challenges in order of <u>most to least</u> problematic when raising broilers. **(Click and drag to rank.)**

______ Necrotic enteritis (254)
______ Airsacculitis (E. *coli* respiratory infection) (255)
______ Gangrenous dermatitis (256)
______ Kinky back (spondylolisthesis) (257)
______ Cellulitis / IP (inflammatory process) (258)
______ Viral enteric disease (259)
______ Bacterial osteomyelitis / lameness (260)
______ Other *(please list)* (261)

---

*Display This Question if NAE = YES CURRENTLY or YES PREVIOUSLY*

Q2.11 Rank these health and welfare challenges in order of most to least problematic when raising broilers without antibiotics.
**(Click and drag to rank.)**

_______ Footpad lesions (357)
_______ Lameness (358)
_______ Corneal lesions (359)
_______ Soiled feathers (360)
_______ Negative impacts on performance (361)
_______ Runting / stunting (362)
_______ Increased morbidity / mortality (363)
_______ Other *(please list)* (364)

---

*Display This Question if NAE = NO*

Q2.12 Rank these health and welfare challenges in order of most to least problematic when raising broilers.
**(Click and drag to rank.)**

_______ Footpad lesions (106)
_______ Lameness (107)
_______ Corneal lesions (108)
_______ Soiled feathers (109)
_______ Negative impacts on performance (110)
_______ Runting / stunting (111)
_______ Increased morbidity / mortality (112)
_______ Other *(please list)* (113)

---

Q2.13 Are there effective tools to prevent or control these disease syndromes without the use antibiotics?

| | Select all that apply | | | | |
|---|---|---|---|---|---|
| | Yes (1) | No (2) | Vaccine (1) | Feed/Water Additive (2) | Management (3) |
| Necrotic enteritis (1) | o | o | o | o | o |
| Airsacculitis (E. *coli* respiratory infection) (2) | o | o | o | o | o |
| Gangrenous dermatitis (3) | o | o | o | o | o |
| Kinky back (spondylolisthesis) (4) | o | o | o | o | o |
| Cellulitis / IP (inflammatory process) (5) | o | o | o | o | o |
| Viral enteric disease (6) | o | o | o | o | o |
| Bacterial osteomyelitis / lameness (7) | o | o | o | o | o |
| Other (please list) (8) | o | o | o | o | o |

---

*Display This Question if NAE = YES CURRENTLY or YES PREVIOUSLY*

Q2.14 Does/did raising broilers without antibiotics impact your production system?

| | Decreased (1) | No change (2) | Increased (3) | Not sure (4) |
|---|---|---|---|---|
| 7-day mortality (1) | o | o | o | o |
| Total mortality (2) | o | o | o | o |
| Feed conversion / average daily gain (3) | o | o | o | o |
| Condemnation (4) | o | o | o | o |

*Display This Question if NAE = NO*

Q2.15 How do you think switching to a raised without antibiotics program would impact your production system?

| | Decreased (1) | No change (2) | Increased (3) | Not sure (4) |
|---|---|---|---|---|
| 7-day mortality (1) | o | o | o | o |
| Total mortality (2) | o | o | o | o |
| Feed conversion / average daily gain (3) | o | o | o | o |
| Condemnation (4) | o | o | o | o |

*Display This Question if NAE = YES CURRENTLY or YES PREVIOUSLY*

Q2.16 Does/did raising broilers without antibiotics necessitate changes in any of the following?

| | Decreased (1) | No change (2) | Increased (3) | Not sure (4) |
|---|---|---|---|---|
| Age at slaughter (1) | o | o | o | o |
| Downtime / layout schedule (2) | o | o | o | o |
| Stocking density (3) | o | o | o | o |
| Grower pay (4) | o | o | o | o |

*Display This Question if NAE = NO*

Q2.17 Do you think switching your animals to a raised without antibiotics program would necessitate changes in any of the following?

| | Decreased (1) | No change (2) | Increased (3) | Not sure (4) |
|---|---|---|---|---|
| Age at slaughter (1) | o | o | o | o |
| Downtime / layout schedule (2) | o | o | o | o |
| Stocking density (3) | o | o | o | o |
| Grower pay (4) | o | o | o | o |

Q2.18 How do you think raised without antibiotics broiler production impacts the following?

| | Significantly improve (11) | Slightly improve (12) | No impact (13) | Slightly worsen (14) | Significantly worsen (15) |
|---|---|---|---|---|---|
| Food safety (15) | o | o | o | o | o |
| Animal health and welfare (16) | o | o | o | o | o |

Q2.19 In your opinion, how do retailers/restaurants/food services think raised without antibiotics broiler production impacts the following?

| | Significantly improve (11) | Slightly improve (12) | No impact (13) | Slightly worsen (14) | Significantly worsen (15) |
|---|---|---|---|---|---|
| Food safety (15) | o | o | o | o | o |
| Animal health and welfare (16) | o | o | o | o | o |

Q2.20 How do you think raised without antibiotics production impacts the cost of broiler production?
- o Significantly increase (1)
- o Slightly increase (2)
- o No impact (3)
- o Slightly decrease (4)
- o Significantly decrease (5)

Q2.21 How do you think raised without antibiotics production impacts the demand for chicken overall by consumers?
- o Significantly increase (1)
- o Slightly increase (2)
- o No impact (3)
- o Slightly decrease (4)
- o Significantly decrease (5)

Q2.22 There are times that maintaining a raised without antibiotics label has priority over flock health and welfare.
- o Strongly agree (1)
- o Somewhat agree (2)
- o Neither agree nor disagree (3)
- o Somewhat disagree (4)
- o Strongly disagree (5)

Q2.23 More stringent health and welfare auditing/assessment is needed for broilers raised without antibiotics.
- o Strongly agree (1)
- o Somewhat agree (2)
- o Neither agree nor disagree (3)
- o Somewhat disagree (4)
- o Strongly disagree (5)

*Display This Question if NAE = YES CURRENTLY or YES PREVIOUSLY*

Q2.24 What is your opinion on the following sentences related to antibiotic use?

| | Strongly agree (1) | Agree (2) | Neutral (3) | Disagree (4) | Strongly disagree (5) | Not sure (6) |
|---|---|---|---|---|---|---|
| Antibiotic use in the broiler industry does not cause problems in humans. (1) | o | o | o | o | o | o |
| Antibiotic use in the broiler industry will make it harder to treat infections in broilers in the future. (2) | o | o | o | o | o | o |
| Antibiotic use in the broiler industry leads to bacterial infections in humans that are more difficult to treat. (3) | o | o | o | o | o | o |
| I would be willing to treat my broilers with antibiotic alternatives if they were equally effective. (4) | o | o | o | o | o | o |

*Display This Question if NAE = NO*

Q2.25 What is your opinion on the following sentences related to antibiotic use?

| | Strongly agree (1) | Agree (2) | Neutral (3) | Disagree (4) | Strongly disagree (5) | Not sure (6) |
|---|---|---|---|---|---|---|
| Antibiotic use in my broilers does not cause problems in humans. (1) | o | o | o | o | o | o |
| Antibiotic use in my broilers will make it harder to treat infections in broilers in the future. (2) | o | o | o | o | o | o |
| Antibiotic use in my broilers leads to bacterial infections in humans that are more difficult to treat. (3) | o | o | o | o | o | o |
| I would be willing to treat my broilers with antibiotic alternatives if they were equally effective. (4) | o | o | o | o | o | o |

Q2.26 What knowledge gaps need to be filled to make a raised without antibiotics production system more successful/sustainable and to have fewer unintended consequences?

________________________________________________________________

________________________________________________________________

________________________________________________________________

________________________________________________________________

________________________________________________________________

**End of Block: Broiler**

### Turkey

Q3.2 Have you ever, past or present, produced/consulted/worked with turkeys enrolled in marketing programs where the animals were raised without the use of any antibiotics?

- o Yes, I am currently working with turkeys being raised without antibiotics
- o Yes, I have previously worked with turkeys being raised without antibiotics but no longer do so
- o No, I have never worked with turkeys raised without antibiotics

*Display This Question if NAE = YES PREVIOUSLY*

Q3.3 Why did you stop working with turkeys raised without the use of any antibiotics?

________________________________________________________________

*Display This Question if NAE = YES CURRENTLY or YES PREVIOUSLY*

Q3.4 Which of these factors contributed to your decision to produce/consult/work with turkeys where the use of antibiotics was not allowed? (Select all that apply)

- o To decrease antibiotic resistance (1)
- o To improve animal health and welfare (2)
- o To increase sale price of animals / product (3)
- o To gain market entry into a retail program (4)
- o To fulfill a client / customer request (5)
- o To eliminate the use of medically important antibiotics (6)
- o Other (please list) (7) ________________________________________________

*Display This Question if NAE = NO*

Q3.5 Which of these factors contributed to your decision not to produce/consult/work with turkeys where the use of antibiotics was not allowed?
(Select all that apply)

- o Not profitable (1)
- o Concerned about negative impacts to animal health and welfare (2)
- o No market pressure (3)
- o Not a sustainable consumer trend (4)
- o Food safety concerns (5)
- o Already eliminated the use of medically important antibiotics (6)
- o Already raising animals in a responsible use program (7)
- o Other (please list) (8) ________________________________________________

(skip to Q3.7)

*Display This Question if NAE = YES CURRENTLY or YES PREVIOUSLY*

Q3.6 Is your experience in producing/consulting/working with turkeys that have been raised without the use of antibiotics part of any of the following types of programs? (Select all that apply)

- o Industry sponsored program (1)
- o Private/Retail/Restaurant/Food Service program (2)
- o Packer/Processor program (3)
- o State/Federal government program (4)
- o Other (please list) (5) ________________________________________________
- o No program (6)

*Display This Question if NAE = NO*

Q3.7 Have you ever considered producing/consulting/working with turkeys raised without the use of antibiotics as part of any of the following types of programs? (Select all that apply)

- o Industry sponsored program (1)
- o Private/Retail/Restaurant/Food Service program (2)
- o Packer/Processor program (3)
- o State/Federal government program (4)
- o Other (please list) (5) ____________________________________________________
- o No program (6)

Q3.8 Are the turkeys you produce/consult/work with currently part of an animal welfare program? (Select all that apply)

- o Industry sponsored quality assurance program (1)
- o Private/Retail/Restaurant/Food Service animal welfare program (2)
- o Packer/Processor animal welfare program (3)
- o State/Federal government animal welfare program (4)
- o Other (please list) (5)
- o No program (6)

*Display This Question if NAE = YES CURRENTLY or YES PREVIOUSLY*

Q3.9 Rank these disease challenges from most to least problematic when raising turkeys without antibiotics.
**(Click and drag to rank.)**

______ Airsacculitis / other respiratory disease (121)
______ Aspergillosis (brooder pneumonia) (122)
______ Cellulitis (123)
______ Coccidiosis (124)
______ Bacterial enteritis (125)
______ Bacterial osteomyelitis / lameness (126)
______ Gangrenous dermatitis (127)
______ Omphalitis (navel infection) (128)
______ Paratyphoid or Salmonellosis (129)
______ Pseudomonas infection (130)
______ Mycoplasmosis (131)
______ Other *(please list)* (132)

*Display This Question if NAE = NO*

Q3.10 Rank these disease challenges in order of most to least problematic when raising turkeys.
**(Click and drag to rank.)**

______ Airsacculitis / other respiratory disease (657)
______ Aspergillosis (brooder pneumonia) (658)
______ Cellulitis (659)
______ Coccidiosis (660)
______ Bacterial enteritis (661)
______ Bacterial osteomyelitis / lameness (662)
______ Gangrenous dermatitis (663)
______ Omphalitis (navel infection) (664)
______ Paratyphoid or Salmonellosis (665)
______ Pseudomonas infection (666)
______ Mycoplasmosis (667)
______ Other *(please list)* (668)

*Display This Question if NAE = YES CURRENTLY or YES PREVIOUSLY*

Q3.11 Rank these health and welfare challenges in order of most to least problematic when raising turkeys without antibiotics.

**(Click and drag to rank.)**

______ Footpad lesions (116)
______ Lameness (117)
______ Corneal lesions (118)
______ Soiled feathers (119)
______ Negative impacts on performance (120)
______ Runting / stunting (121)
______ Increased morbidity / mortality (122)
______ Other *(please list)* (123)

---

*Display This Question if NAE = NO*

Q3.12 Rank these health and welfare challenges in order of most to least problematic when raising turkeys.

**(Click and drag to rank.)**

______ Footpad lesions (102)
______ Lameness (103)
______ Corneal lesions (104)
______ Soiled feathers (105)
______ Negative impacts on performance (106)
______ Runting / stunting (107)
______ Increased morbidity / mortality (108)
______ Other *(please list)* (109)

---

Q3.13 Are there effective tools to prevent or control these disease syndromes without the use antibiotics?

| | Select all that apply | | | | |
|---|---|---|---|---|---|
| | Yes (1) | No (2) | Vaccine (1) | Feed/Water Additive (2) | Management (3) |
| Airsacculitis / other respiratory disease (1) | o | o | o | o | o |
| Aspergillosis (brooder pneumonia) (2) | o | o | o | o | o |
| Cellulitis (3) | o | o | o | o | o |
| Coccidiosis (4) | o | o | o | o | o |
| Bacterial enteritits (5) | o | o | o | o | o |
| Bacterial osteomyelitis / lameness (6) | o | o | o | o | o |
| Gangrenous dermatitis (7) | o | o | o | o | o |
| Omphalitis (navel infection) (8) | o | o | o | o | o |
| Paratyphoid or Salmonellosis (9) | o | o | o | o | o |
| Pseudomonas infection (10) | o | o | o | o | o |
| Mycoplasmosis (11) | o | o | o | o | o |
| Other (please list) (12) | o | o | o | o | o |

*Display This Question if NAE = YES CURRENTLY or YES PREVIOUSLY*

Q3.14 Does/did raising turkeys without antibiotics impact your production system?

| | Decreased (1) | No change (2) | Increased (3) | Not sure (4) |
|---|---|---|---|---|
| 7-day mortality (1) | o | o | o | o |
| Total mortality (livability) (2) | o | o | o | o |
| Feed conversion / average daily gain (3) | o | o | o | o |
| Condemnation (4) | o | o | o | o |

*Display This Question if NAE = NO*

Q3.15 How do you think switching to a raised without antibiotics program would impact your production system?

| | Decreased (1) | No change (2) | Increased (3) | Not sure (4) |
|---|---|---|---|---|
| 7-day mortality (1) | o | o | o | o |
| Total mortality (livability) (2) | o | o | o | o |
| Feed conversion / average daily gain (3) | o | o | o | o |
| Condemnation (4) | o | o | o | o |

*Display This Question if NAE = YES CURRENTLY or YES PREVIOUSLY*

Q3.16 Does/did raising turkeys without antibiotics necessitate changes in any of the following?

| | Decreased (1) | No change (2) | Increased (3) | Not sure (4) |
|---|---|---|---|---|
| Age at slaughter (1) | o | o | o | o |
| Downtime / layout schedule (2) | o | o | o | o |
| Stocking density (3) | o | o | o | o |
| Grower pay (4) | o | o | o | o |

*Display This Question if NAE = NO*

Q3.17 Do you think switching your animals to a raised without antibiotics program would necessitate changes in any of the following?

| | Decreased (1) | No change (2) | Increased (3) | Not sure (4) |
|---|---|---|---|---|
| Age at slaughter (1) | o | o | o | o |
| Downtime / layout schedule (2) | o | o | o | o |
| Stocking density (3) | o | o | o | o |
| Grower pay (4) | o | o | o | o |

Q3.18 How do you think raised without antibiotics turkey production impacts the following?

| | Significantly improve (11) | Slightly improve (12) | No impact (13) | Slightly worsen (14) | Significantly worsen (15) |
|---|---|---|---|---|---|
| Food safety (15) | o | o | o | o | o |
| Animal health and welfare (16) | o | o | o | o | o |

Q3.19 In your opinion, how do retailers/restaurants/food services think raised without antibiotics turkey production impacts the following?

| | Significantly improve (11) | Slightly improve (12) | No impact (13) | Slightly worsen (14) | Significantly worsen (15) |
|---|---|---|---|---|---|
| Food safety (15) | o | o | o | o | o |
| Animal health and welfare (16) | o | o | o | o | o |

Q3.20 How do you think raised without antibiotics production impacts the cost of turkey production?

- o Significantly increase (1)
- o Slightly increase (2)
- o No impact (3)
- o Slightly decrease (4)
- o Significantly decrease (5)

Q3.21 How do you think raised without antibiotics production impacts the demand for turkey overall by consumers?

- o Significantly increase (1)
- o Slightly increase (2)
- o No impact (3)
- o Slightly decrease (4)
- o Significantly decrease (5)

Q3.22 There are times that maintaining a raised without antibiotics label has priority over flock health and welfare.

- o Strongly agree (1)
- o Somewhat agree (2)
- o Neither agree nor disagree (3)
- o Somewhat disagree (4)
- o Strongly disagree (5)

Q3.23 More stringent health and welfare auditing/assessment is needed for turkeys raised without antibiotics.

- o Strongly agree (1)
- o Somewhat agree (2)
- o Neither agree nor disagree (3)
- o Somewhat disagree (4)
- o Strongly disagree (5)

*Display This Question if NAE = YES CURRENTLY or YES PREVIOUSLY*

Q3.24 What is your opinion on the following sentences related to antibiotic use?

| | Strongly agree (1) | Agree (2) | Neutral (3) | Disagree (4) | Strongly disagree (5) | Not sure (6) |
|---|---|---|---|---|---|---|
| Antibiotic use in the turkey industry does not cause problems in humans. (1) | o | o | o | o | o | o |
| Antibiotic use in the turkey industry will make it harder to treat infections in turkeys in the future. (2) | o | o | o | o | o | o |
| Antibiotic use in the turkey industry leads to bacterial infections in humans that are more difficult to treat. (3) | o | o | o | o | o | o |
| I would be willing to treat my turkeys with antibiotic alternatives if they were equally effective. (4) | o | o | o | o | o | o |

*Display This Question if NAE = NO*

Q3.25 What is your opinion on the following sentences related to antibiotic use?

| | Strongly agree (1) | Agree (2) | Neutral (3) | Disagree (4) | Strongly disagree (5) | Not sure (6) |
|---|---|---|---|---|---|---|
| Antibiotic use in my turkeys does not cause problems in humans. (1) | o | o | o | o | o | o |
| Antibiotic use in my turkeys will make it harder to treat infections in turkeys in the future. (2) | o | o | o | o | o | o |
| Antibiotic use in my turkeys leads to bacterial infections in humans that are more difficult to treat. (3) | o | o | o | o | o | o |
| I would be willing to treat my turkeys with antibiotic alternatives if they were equally effective. (4) | o | o | o | o | o | o |

Q3.26 What knowledge gaps need to be filled to make a raised without antibiotics production system more successful/sustainable and to have fewer unintended consequences?

________________________________________________________________

________________________________________________________________

________________________________________________________________

________________________________________________________________

________________________________________________________________

**End of Block: Turkey**

### Swine

Q4.2 Have you ever, past or present, produced/consulted/worked with swine enrolled in marketing programs where the animals were raised without the use of any antibiotics?

- Yes, I am currently working with swine being raised without antibiotics
- Yes, I have previously worked with swine being raised without antibiotics but no longer do so
- No, I have never worked with swine raised without antibiotics

*Display This Question if NAE = YES PREVIOUSLY*

Q4.3 Why did you stop working with swine raised without the use of any antibiotics?

________________________________________________________________

---

*Display This Question if NAE = YES CURRENTLY or YES PREVIOUSLY*

Q4.4 Which of these factors contributed to your decision to produce/consult/work with swine where the use of antibiotics was not allowed? (Select all that apply)

- To decrease antibiotic resistance (1)
- To improve animal health and welfare (2)
- To increase sale price of animals/product (3)
- To gain market entry into a retail program (4)
- To fulfill a client/customer request (5)
- To eliminate the use of medically important antibiotics (6)
- Other (please list) (7) ________________________________________________

---

*Display This Question if NAE = NO*

Q4.5 Which of these factors contributed to your decision not to produce/consult/work with swine where the use of antibiotics was not allowed?
(Select all that apply)

- Not profitable (1)
- Concerned about negative impacts to animal health and welfare (2)
- No market pressure (3)
- Not a sustainable consumer trend (4)
- Food safety concerns (5)
- Already eliminated the use of medically important antibiotics (6)
- Already raising animals in a responsible use program (7)
- Other (please list) (8) ________________________________________________

---

*Display This Question if NAE = YES CURRENTLY or YES PREVIOUSLY*

Q4.6 Is your experience in producing/consulting/working with swine that have been raised without the use of antibiotics part of any of the following types of programs? (Select all that apply)

- Industry sponsored program (1)
- Private/Retail/Restaurant/Food Service program (2)
- Packer/Processor program (3)
- State/Federal government program (4)
- Other (please list) (5) ________________________________________________
- No program (6)

---

*Display This Question if NAE = NO*

Q4.7 Have you ever considered producing/consulting/working with swine raised without the use of antibiotics as part of any of the following types of programs? (Select all that apply)

- o Industry sponsored program (1)
- o Private/Retail/Restaurant/Food Service program (2)
- o Packer/Processor program (3)
- o State/Federal government program (4)
- o Other (please list) (5) ____________________________________________________
- o No program (6)

Q4.8 Are the swine you produce/consult/work with currently part of an animal welfare program? (Select all that apply)

- o PQA Plus/Common Swine Industry Audit (1)
- o NOS – National Organic Standard (2)
- o GAP (3)
- o Certified Humane ( Humane Farm Animal Care) (4)
- o American Humane Certified (5)
- o Animal Welfare Approved (6)
- o Privately owned/facilitated animal welfare program (7)
- o No program (8)

*Display This Question if NAE = YES CURRENTLY or YES PREVIOUSLY*

Q4.9 Rank these disease challenges from most to least problematic when raising swine without antibiotics.
**(Click and drag to rank.)**

______ Post weaning hemolytic *E. coli* (254)
______ Ileitis (255)
______ Mycoplasma pneumonia (256)
______ *Actinobacillus suis*, *Haemophilus parasuis* and *Streptococcus suis* (257)
______ Salmonella (258)
______ Erysipelas (259)
______ Other *(please list)* (260)

*Display This Question if NAE = NO*

Q4.10 Rank these disease challenges in order of most to least problematic when raising swine.
**(Click and drag to rank.)**

______ Post weaning hemolytic *E. coli* (254)
______ Ileitis (255)
______ Mycoplasma pneumonia (256)
______ *Actinobacillus suis*, *Haemophilus parasuis* and *Streptococcus suis* (257)
______ Salmonella (258)
______ Erysipelas (259)
______ Other *(please list)* (260)

*Display This Question if NAE = YES CURRENTLY or YES PREVIOUSLY*

Q4.11 Rank these health and welfare challenges in order of most to least problematic when raising swine without antibiotics.
**(Click and drag to rank.)**

______ Respiratory system disorders (433)
______ Musculoskeletal system disorders (434)
______ Central nervous center disorders (435)
______ Digestive system disorders (436)
______ Reproductive system disorders (437)
______ Damaging behaviors (tail/ear/flank biting) (438)
______ Other *(please list)* (439)

*Display This Question if NAE = NO*

Q4.12 Rank these health and welfare challenges in order of most to least problematic when raising swine.
**(Click and drag to rank.)**

______ Respiratory system disorders (129)
______ Musculoskeletal system disorders (130)
______ Central nervous center disorders (131)
______ Digestive system disorders (132)
______ Reproductive system disorders (133)
______ Damaging behaviors (tail/ear/flank biting) (134)
______ Other *(please list)* (135)

Q4.13 Are there effective tools to prevent or control these disease syndromes without the use antibiotics?

| | Select all that apply | | | | |
|---|---|---|---|---|---|
| | Yes (1) | No (2) | Vaccine (1) | Feed/Water Additive (2) | Management (3) |
| Post weaning hemolytic *E. coli* (1) | o | o | o | o | o |
| Ileitis (2) | o | o | o | o | o |
| Mycoplasma pneumonia (3) | o | o | o | o | o |
| *Actinobacillus suis, Haemophilus parasuis* and *Streptococcus suis* (4) | o | o | o | o | o |
| Salmonella (5) | o | o | o | o | o |
| Erysipelas (6) | o | o | o | o | o |
| Other (please list) (7) | o | o | o | o | o |

*Display This Question if NAE = YES CURRENTLY or YES PREVIOUSLY*

Q4.14 Does/did raising swine without antibiotics necessitate changes in any of these management strategies or facility designs?

| | Yes (1) | No (2) | Not sure (3) |
|---|---|---|---|
| Weaning age (1) | o | o | o |
| Air filtering (2) | o | o | o |
| Personnel requirements (3) | o | o | o |
| Biosecurity changes (cleaning, disinfection, visitor policy, etc.) (4) | o | o | o |
| Space allowance (5) | o | o | o |
| Bedding (6) | o | o | o |
| Other (please list) (7) | o | o | o |

---

*Display This Question if NAE = NO*

Q4.15 Do you think switching to a raised without antibiotics program would necessitate changes in any of these management strategies or facility designs?

| | Yes (1) | No (2) | Not sure (3) |
|---|---|---|---|
| Weaning age (1) | o | o | o |
| Air filtering (2) | o | o | o |
| Personnel requirements (3) | o | o | o |
| Biosecurity changes (cleaning, disinfection, visitor policy, etc.) (4) | o | o | o |
| Space allowance (5) | o | o | o |
| Bedding (6) | o | o | o |
| Other (please list) (7) | o | o | o |

---

*Display This Question if NAE = YES CURRENTLY or YES PREVIOUSLY*

Q4.16 Does/did raising swine without antibiotics impact your production system?

| | Decreased (1) | No change (2) | Increased (3) | Not sure (4) |
|---|---|---|---|---|
| Feed efficiency (1) | o | o | o | o |
| Morbidity (2) | o | o | o | o |
| Mortality (3) | o | o | o | o |
| Age at slaughter (4) | o | o | o | o |
| Weight at slaughter (5) | o | o | o | o |

---

*Display This Question if NAE = NO*

Q4.17 How do you think switching to a raised without antibiotics program would impact your production system?

| | Decreased (1) | No change (2) | Increased (3) | Not sure (4) |
|---|---|---|---|---|
| Feed efficiency (1) | o | o | o | o |
| Morbidity (2) | o | o | o | o |
| Mortality (3) | o | o | o | o |
| Age at slaughter (4) | o | o | o | o |
| Weight at slaughter (5) | o | o | o | o |

Q4.18 How do you think raised without antibiotics swine production impacts the following?

| | Significantly improve (11) | Slightly improve (12) | No impact (13) | Slightly worsen (14) | Significantly worsen (15) |
|---|---|---|---|---|---|
| Food safety (15) | o | o | o | o | o |
| Animal health and welfare (16) | o | o | o | o | o |

Q4.19 In your opinion, how do retailers/restaurants/food services think raised without antibiotics swine production impacts the following?

| | Significantly improve (11) | Slightly improve (12) | No impact (13) | Slightly worsen (14) | Significantly worsen (15) |
|---|---|---|---|---|---|
| Food safety (15) | o | o | o | o | o |
| Animal health and welfare (16) | o | o | o | o | o |

Q4.20 How do you think raised without antibiotics production impacts the cost of swine production?

- o Significantly increase (1)
- o Slightly increase (2)
- o No impact (3)
- o Slightly decrease (4)
- o Significantly decrease (5)

Q4.21 How do you think raised without antibiotics production impacts the demand for pork overall by consumers?
- Significantly increase (1)
- Slightly increase (2)
- No impact (3)
- Slightly decrease (4)
- Significantly decrease (5)

Q4.22 There are times that maintaining a raised without antibiotics label has priority over herd health and welfare.
- Strongly agree (1)
- Somewhat agree (2)
- Neither agree nor disagree (3)
- Somewhat disagree (4)
- Strongly disagree (5)

Q4.23 More stringent health and welfare auditing/assessment is needed for swine raised without antibiotics.
- Strongly agree (1)
- Somewhat agree (2)
- Neither agree nor disagree (3)
- Somewhat disagree (4)
- Strongly disagree (5)

*Display This Question if NAE = YES CURRENTLY or YES PREVIOUSLY*

Q4.24 What is your opinion on the following sentences related to antibiotic use?

| | Strongly agree (1) | Agree (2) | Neutral (3) | Disagree (4) | Strongly disagree (5) | Not sure (6) |
|---|---|---|---|---|---|---|
| Antibiotic use in the swine industry does not cause problems in humans. (1) | o | o | o | o | o | o |
| Antibiotic use in the swine industry will make it harder to treat infections in swine in the future. (2) | o | o | o | o | o | o |
| Antibiotic use in the swine industry leads to bacterial infections in humans that are more difficult to treat. (3) | o | o | o | o | o | o |
| I would be willing to treat my swine with antibiotic alternatives if they were equally effective. (4) | o | o | o | o | o | o |

*Display This Question if NAE = NO*

Q4.25 What is your opinion on the following sentences related to antibiotic use?

| | Strongly agree (1) | Agree (2) | Neutral (3) | Disagree (4) | Strongly disagree (5) | Not sure (6) |
|---|---|---|---|---|---|---|
| Antibiotic use in my swine does not cause problems in humans. (1) | o | o | o | o | o | o |
| Antibiotic use in my swine will make it harder to treat infections in swine in the future. (2) | o | o | o | o | o | o |
| Antibiotic use in my swine leads to bacterial infections in humans that are more difficult to treat. (3) | o | o | o | o | o | o |
| I would be willing to treat my swine with antibiotic alternatives if they were equally effective. (4) | o | o | o | o | o | o |

Q4.26 What knowledge gaps need to be filled to make a raised without antibiotics production system more successful/sustainable and to have fewer unintended consequences?

________________________________________________________________

________________________________________________________________

________________________________________________________________

________________________________________________________________

________________________________________________________________

**End of Block: Swine**

### Beef

Q5.2 Have you ever, past or present, produced/consulted/worked with beef cattle enrolled in marketing programs where the animals were raised without the use of any antibiotics?

- o Yes, I am currently working with beef cattle being raised without antibiotics
- o Yes, I have previously worked with beef cattle being raised without antibiotics but no longer do so
- o No, I have never worked with beef cattle raised without antibiotics

*Display This Question if NAE = YES PREVIOUSLY*

Q5.3 Why did you stop working with beef cattle raised without the use of any antibiotics?

________________________________________________________________

*Display This Question if NAE = YES CURRENTLY or YES PREVIOUSLY*

Q5.4 Which of these factors contributed to your decision to produce/consult/work with beef cattle where the use of antibiotics was not allowed? (Select all that apply)

- o To decrease antibiotic resistance (1)
- o To improve animal health and welfare (2)
- o To increase sale price of animals/product (3)
- o To gain market entry into a retail program (4)
- o To fulfill a client/customer request (5)
- o To eliminate the use of medically important antibiotics (6)
- o Other (please list) (7) ________________________________________________

*Display This Question if NAE = NO*

Q5.5 Which of these factors contributed to your decision <u>not</u> to produce/consult/work with beef cattle where the use of antibiotics was not allowed?
(Select all that apply)

- o Not profitable (1)
- o Concerned about negative impacts to animal health and welfare (2)
- o No market pressure (3)
- o Not a sustainable consumer trend (4)
- o Food safety concerns (5)
- o Already eliminated the use of medically important antibiotics (6)
- o Already raising animals in a responsible use program (7)
- o Other (please list) (8) ________________________________________________

*Display This Question if NAE = YES CURRENTLY or YES PREVIOUSLY*

Q5.6 Is your experience in producing/consulting/working with beef cattle that have been raised without the use of antibiotics part of any of the following types of programs? (Select all that apply)

- o Industry sponsored program (1)
- o Private/Retail/Restaurant/Food Service program (2)
- o Packer/Processor program (3)
- o State/Federal government program (4)
- o Other (please list) (5) ________________________________________________
- o No program (6)

*Display This Question if NAE = NO*

Q5.7 Have you ever considered producing/consulting/working with beef cattle raised without the use of antibiotics as part of any of the following types of programs? (Select all that apply)

- Industry sponsored program (1)
- Private/Retail/Restaurant/Food Service program (2)
- Packer/Processor program (3)
- State/Federal government program (4)
- Other (please list) (5) ____________________
- No program (6)

---

Q5.8 Are the beef cattle you produce/consult/work with currently part of an animal welfare program? (Select all that apply)

- Industry sponsored quality assurance program (1)
- Private/Retail/Restaurant/Food Service animal welfare program (2)
- Packer/Processor animal welfare program (3)
- State/Federal government animal welfare program (4)
- Other (please list) (5) ____________________
- No program (6)

---

*Display This Question if NAE = YES CURRENTLY or YES PREVIOUSLY*

Q5.9 Rank these disease challenges from most to least problematic when raising beef cattle without antibiotics.
**(Click and drag to rank.)**

______ Calf scours (306)
______ Pink eye (307)
______ Anaplasmosis (308)
______ Foot rot lameness (309)
______ Digital Dermatitis (310)
______ Bovine Respiratory Disease (311)
______ Liver abscessation (312)
______ Central nervous system disorders (313)
______ Other *(please list)* (314)

---

*Display This Question if NAE = NO*

Q5.10 Rank these disease challenges in order of most to least problematic when raising beef cattle.
**(Click and drag to rank.)**

______ Calf scours (254)
______ Pink eye (255)
______ Anaplasmosis (256)
______ Foot rot lameness (257)
______ Digital Dermatitis (258)
______ Bovine Respiratory Disease (259)
______ Liver abscessation (260)
______ Central nervous system disorders (261)
______ Other *(please list)* (262)

---

*Display This Question if NAE = YES CURRENTLY or YES PREVIOUSLY*

Q5.11 Rank these health and welfare challenges in order of most to least problematic when raising beef cattle without antibiotics.
**(Click and drag to rank.)**

______ Respiratory system disorders (212)
______ Musculoskeletal system disorders (213)
______ Central nervous system disorders (214)
______ Digestive system disorders (215)
______ Reproductive system disorders (216)
______ Other *(please list)* (217)

---

*Display This Question if NAE = NO*

Q5.12 Rank these health and welfare challenges in order of most to least problematic when raising beef cattle.
**(Click and drag to rank.)**

______ Respiratory system disorders (107)
______ Musculoskeletal system disorders (108)
______ Central nervous system disorders (109)
______ Digestive system disorders (110)
______ Reproductive system disorders (111)
______ Other *(please list)* (112)

---

Q5.13 Are there effective tools to prevent or control these disease syndromes without the use antibiotics?

| | Select all that apply | | | | |
|---|---|---|---|---|---|
| | Yes (1) | No (2) | Vaccine (1) | Feed/Water Additive (2) | Management (3) |
| Calf scours (1) | o | o | o | o | o |
| Pink eye (2) | o | o | o | o | o |
| Anaplasmosis (3) | o | o | o | o | o |
| Foot rot lameness (4) | o | o | o | o | o |
| Digital Dermatitis (5) | o | o | o | o | o |
| Bovine Respiratory Disease (6) | o | o | o | o | o |
| Liver abscessation (7) | o | o | o | o | o |
| Central nervous system disorders (8) | o | o | o | o | o |
| Other (please list) (9) | o | o | o | o | o |

---

*Display This Question if NAE = YES CURRENTLY or YES PREVIOUSLY*

Q5.14 Does/did raising beef cattle without antibiotics impact your production system?

| | Decreased (1) | No change (2) | Increased (3) | Not sure (4) |
|---|---|---|---|---|
| Personnel requirements (1) | o | o | o | o |
| Feed efficiency (2) | o | o | o | o |
| Cattle morbidity (3) | o | o | o | o |
| Death loss (4) | o | o | o | o |
| Age at slaughter (5) | o | o | o | o |

*Display This Question if NAE = NO*

Q5.15 How do you think switching to a raised without antibiotics program would impact your production system?

| | Decreased (1) | No change (2) | Increased (3) | Not sure (4) |
|---|---|---|---|---|
| Personnel requirements (1) | o | o | o | o |
| Feed efficiency (2) | o | o | o | o |
| Cattle morbidity (3) | o | o | o | o |
| Death loss (4) | o | o | o | o |
| Age at slaughter (5) | o | o | o | o |

Q5.16 How do <u>you</u> think raised without antibiotics beef cattle production impacts the following?

| | Significantly improve (11) | Slightly improve (12) | No impact (13) | Slightly worsen (14) | Significantly worsen (15) |
|---|---|---|---|---|---|
| Food safety (15) | o | o | o | o | o |
| Animal health and welfare (16) | o | o | o | o | o |

Q5.17 In your opinion, how do retailers/restaurants/food services think raised without antibiotics beef cattle production impacts the following?

| | Significantly improve (11) | Slightly improve (12) | No impact (13) | Slightly worsen (14) | Significantly worsen (15) |
|---|---|---|---|---|---|
| Food safety (15) | o | o | o | o | o |
| Animal health and welfare (16) | o | o | o | o | o |

Q5.18 How do you think raised without antibiotics production impacts the cost of beef cattle production?
- o Significantly increase (1)
- o Slightly increase (2)
- o No impact (3)
- o Slightly decrease (4)
- o Significantly decrease (5)

Q5.19 How do you think raised without antibiotics production impacts the demand for beef overall by consumers?
- o Significantly increase (1)
- o Slightly increase (2)
- o No impact (3)
- o Slightly decrease (4)
- o Significantly decrease (5)

Q5.20 There are times that maintaining a raised without antibiotics label has priority over herd health and welfare.
- o Strongly agree (1)
- o Somewhat agree (2)
- o Neither agree nor disagree (3)
- o Somewhat disagree (4)
- o Strongly disagree (5)

Q5.21 More stringent health and welfare auditing/assessment is needed for beef cattle raised without antibiotics.
- o Strongly agree (1)
- o Somewhat agree (2)
- o Neither agree nor disagree (3)
- o Somewhat disagree (4)
- o Strongly disagree (5)

*Display This Question if NAE = YES CURRENTLY or YES PREVIOUSLY*

Q5.22 What is your opinion on the following sentences related to antibiotic use?

| | Strongly agree (1) | Agree (2) | Neutral (3) | Disagree (4) | Strongly disagree (5) | Not sure (6) |
|---|---|---|---|---|---|---|
| Antibiotic use in the beef cattle industry does not cause problems in humans. (1) | o | o | o | o | o | o |
| Antibiotic use in beef cattle industry will make it harder to treat infections in beef cattle in the future. (2) | o | o | o | o | o | o |
| Antibiotic use in beef cattle industry leads to bacterial infections in humans that are more difficult to treat. (3) | o | o | o | o | o | o |
| I would be willing to treat my beef cattle with antibiotic alternatives if they were equally effective. (4) | o | o | o | o | o | o |

*Display This Question if NAE = NO*

Q5.23 What is your opinion on the following sentences related to antibiotic use?

| | Strongly agree (1) | Agree (2) | Neutral (3) | Disagree (4) | Strongly disagree (5) | Not sure (6) |
|---|---|---|---|---|---|---|
| Antibiotic use in my beef cattle does not cause problems in humans. (1) | o | o | o | o | o | o |
| Antibiotic use in my beef cattle will make it harder to treat infections in beef cattle in the future. (2) | o | o | o | o | o | o |
| Antibiotic use in my beef cattle leads to bacterial infections in humans that are more difficult to treat. (3) | o | o | o | o | o | o |
| I would be willing to treat my beef cattle with antibiotic alternatives if they were equally effective. (4) | o | o | o | o | o | o |

Q5.24 What knowledge gaps need to be filled to make a raised without antibiotics production system more successful/sustainable and to have fewer unintended consequences?

________________________________________________________________

________________________________________________________________

________________________________________________________________

________________________________________________________________

**End of Block: Beef**

### Dairy

Q6.2 Have you ever, past or present, produced/consulted/worked with dairy cattle/calves enrolled in marketing programs where the animals were raised without the use of any antibiotics?
- o Yes, I am currently working with dairy cattle/calves being raised without antibiotics (3)
- o Yes, I have previously worked with dairy cattle/calves being raised without antibiotics but no longer do so (4)
- o No, I have never worked with dairy cattle/calves raised without antibiotics (5)

---

*Display This Question if NAE = YES PREVIOUSLY*

Q6.3 Why did you stop working with dairy cattle/calves raised without the use of any antibiotics?

________________________________________________________________

---

*Display This Question if NAE = YES CURRENTLY or YES PREVIOUSLY*

Q6.4 Which of these factors contributed to your decision to produce/consult/work with dairy cattle/calves where the use of antibiotics was not allowed? (Select all that apply)
- o To decrease antibiotic resistance (1)
- o To improve animal health and welfare (2)
- o To increase sale price of animals/product (3)
- o To gain market entry into a retail program (4)
- o To fulfill a client/customer request (5)
- o To eliminate the use of medically important antibiotics (6)
- o Other (please list) (7) ________________________________________________

*Display This Question if NAE = NO*

Q6.5 Which of these factors contributed to your decision <u>not</u> to produce/consult/work with dairy cattle/calves where the use of antibiotics was not allowed?
(Select all that apply)
- o Not profitable (1)
- o Concerned about negative impacts to animal health and welfare (2)
- o No market pressure (3)
- o Not a sustainable consumer trend (4)
- o Food safety concerns (5)
- o Already eliminated the use of medically important antibiotics (6)
- o Already raising animals in a responsible use program (7)
- o Other (please list) (8) ________________________________________________

---

*Display This Question if NAE = YES CURRENTLY or YES PREVIOUSLY*

Q6.6 Is your experience in producing/consulting/working with dairy cattle/calves that have been raised without the use of antibiotics part of any of the following types of programs? (Select all that apply)
- o Industry sponsored program (1)
- o Private/Retail/Restaurant/Food Service program (2)
- o Packer/Processor program (3)
- o State/Federal government program (4)
- o Other (please list) (5) ________________________________________________
- o No program (6)

---

*Display This Question if NAE = NO*

Q6.7 Have you ever considered producing/consulting/working with dairy cattle/calves raised without the use of antibiotics as part of any of the following types of programs? (Select all that apply)

- o Industry sponsored program (1)
- o Private/Retail/Restaurant/Food Service program (2)
- o Packer/Processor program (3)
- o State/Federal government program (4)
- o Other (please list) (5) ____________________________________________________
- o No program (6)

---

Q6.8 Are the dairy cattle/calves you produce/consult/work with currently part of an animal welfare program? (Select all that apply)

- o Industry sponsored quality assurance program (e.g. FARM) (1)
- o Private/Retail/Restaurant/Food Service animal welfare program (2)
- o Packer/Processor animal welfare program (3)
- o State/Federal government animal welfare program (4)
- o Other (please list) (5) ____________________________________________________
- o No program (6)

---

*Display This Question if NAE = YES CURRENTLY or YES PREVIOUSLY*

Q6.9 Rank these disease challenges from most to least problematic when raising dairy cattle/calves without antibiotics.

**(Click and drag to rank.)**

______ Calf scours (134)
______ Metritis (135)
______ Mastitis (136)
______ Anaplasmosis (137)
______ Metabolic disorder (i.e. ketosis, milk fever) (138)
______ Foot rot lameness (139)
______ Digital Dermititis (140)
______ Bovine Respiratory Disease (141)
______ Other *(please list)* (143)

---

*Display This Question if NAE = NO*

Q6.10 Rank these disease challenges in order of most to least problematic when raising dairy cattle/calves.

**(Click and drag to rank.)**

______ Calf scours (262)
______ Metritis (263)
______ Mastitis (264)
______ Anaplasmosis (265)
______ Metabolic disorder (i.e. ketosis, milk fever) (266)
______ Foot rot lameness (267)
______ Digital Dermititis (268)
______ Bovine Respiratory Disease (269)
______ Other *(please list)* (270)

---

*Display This Question if NAE = YES CURRENTLY or YES PREVIOUSLY*

Q6.11 Rank these health and welfare challenges in order of most to least problematic when raising dairy cattle/calves without antibiotics.

**(Click and drag to rank.)**

______ Mammary gland disorders (126)
______ Respiratory system disorders (127)
______ Pneumonia (128)
______ Musculoskeletal system disorders (129)
______ Central nervous center disorders (130)
______ Digestive system disorders (131)
______ Reproductive system disorders (132)
______ Other *(please list)* (133)

*Display This Question if NAE = NO*

Q6.12 Rank these health and welfare challenges in order of most to least problematic when raising dairy cattle/calves.

**(Click and drag to rank.)**

______ Mammary gland disorders (114)
______ Respiratory system disorders (115)
______ Pneumonia (116)
______ Musculoskeletal system disorders (117)
______ Central nervous center disorders (118)
______ Digestive system disorders (119)
______ Reproductive system disorders (120)
______ Other *(please list)* (121)

Q6.13 Are there effective tools to prevent or control these disease syndromes without the use antibiotics?

| | Select all that apply | | | | |
|---|---|---|---|---|---|
| | Yes (1) | No (2) | Vaccine (1) | Feed/Water Additive (2) | Management (3) |
| Calf scours (1) | o | o | o | o | o |
| Metritis (2) | o | o | o | o | o |
| Mastitis (3) | o | o | o | o | o |
| Anaplasmosis (4) | o | o | o | o | o |
| Metabolic disorder (i.e. ketosis, milk fever) (5) | o | o | o | o | o |
| Foot rot lameness (6) | o | o | o | o | o |
| Digital Dermititis (7) | o | o | o | o | o |
| Bovine Respiratory Disease (8) | o | o | o | o | o |
| Other (please list) (9) | o | o | o | o | o |

*Display This Question if NAE = YES CURRENTLY or YES PREVIOUSLY*

Q6.14 Does/did raising dairy cattle/calves without antibiotics impact your production system?

| | Decreased (1) | No change (2) | Increased (3) | Not sure (4) |
|---|---|---|---|---|
| Number of personnel required (1) | o | o | o | o |
| Feed efficiency (2) | o | o | o | o |
| Milk yield performance (3) | o | o | o | o |
| Culling rate (4) | o | o | o | o |
| Cattle morbidity (5) | o | o | o | o |
| Somatic cell count (6) | o | o | o | o |

*Display This Question if NAE = NO*

Q6.15 How do you think switching to a raised without antibiotics program would impact your production system?

| | Decreased (1) | No change (2) | Increased (3) | Not sure (4) |
|---|---|---|---|---|
| Number of personnel required (1) | o | o | o | o |
| Feed efficiency (2) | o | o | o | o |
| Milk yield performance (3) | o | o | o | o |
| Culling rate (4) | o | o | o | o |
| Cattle morbidity (5) | o | o | o | o |
| Somatic cell count (6) | o | o | o | o |

Q6.16 How do <u>you</u> think raised without antibiotics dairy cattle/calf production impacts the following?

| | Significantly improve (11) | Slightly improve (12) | No impact (13) | Slightly worsen (14) | Significantly worsen (15) |
|---|---|---|---|---|---|
| Food safety (15) | o | o | o | o | o |
| Animal health and welfare (16) | o | o | o | o | o |

Q6.17 In your opinion, how do retailers/restaurants/food services think raised without antibiotics dairy cattle/calves production impacts the following?

| | Significantly improve (11) | Slightly improve (12) | No impact (13) | Slightly worsen (14) | Significantly worsen (15) |
|---|---|---|---|---|---|
| Food safety (15) | o | o | o | o | o |
| Animal health and welfare (16) | o | o | o | o | o |

Q6.18 How do you think raised without antibiotics production impacts the cost of dairy cattle/calf production?
- o Significantly increase (1)
- o Slightly increase (2)
- o No impact (3)
- o Slightly decrease (4)
- o Significantly decrease (5)

Q6.19 How do you think raised without antibiotics production impacts the demand for milk and dairy products overall by consumers?
- o Significantly increase (1)
- o Slightly increase (2)
- o No impact (3)
- o Slightly decrease (4)
- o Significantly decrease (5)

Q6.20 There are times that maintaining a raised without antibiotics label has priority over herd health and welfare.
- o Strongly agree (1)
- o Agree (2)
- o Somewhat agree (3)
- o Neither agree nor disagree (4)
- o Somewhat disagree (5)
- o Disagree (6)

Q6.21 More stringent health and welfare auditing/assessment is needed for dairy cattle/calves raised without antibiotics.
- o Strongly agree (1)
- o Somewhat agree (2)
- o Neither agree nor disagree (3)
- o Somewhat disagree (4)
- o Strongly disagree (5)

*Display This Question if NAE = YES CURRENTLY or YES PREVIOUSLY*

Q6.22 What is your opinion on the following sentences related to antibiotic use?

| | Strongly agree (1) | Agree (2) | Neutral (3) | Disagree (4) | Strongly disagree (5) | Not sure (6) |
|---|---|---|---|---|---|---|
| Antibiotic use in the dairy industry does not cause problems in humans. (1) | o | o | o | o | o | o |
| Antibiotic use in the dairy industry will make it harder to treat infections in dairy cattle/calves in the future. (2) | o | o | o | o | o | o |
| Antibiotic use in dairy industry leads to bacterial infections in humans that are more difficult to treat. (3) | o | o | o | o | o | o |
| There are effective alternatives to antibiotics for me to treat my dairy cattle/calves. (4) | o | o | o | o | o | o |

*Display This Question if NAE = NO*

Q6.23 What is your opinion on the following sentences related to antibiotic use?

| | Strongly agree (1) | Agree (2) | Neutral (3) | Disagree (4) | Strongly disagree (5) | Not sure (6) |
|---|---|---|---|---|---|---|
| Antibiotic use in my dairy cattle/calves does not cause problems in humans. (1) | o | o | o | o | o | o |
| Antibiotic use in my dairy cattle/calves will make it harder to treat infections in dairy cattle/calves in the future. (2) | o | o | o | o | o | o |
| Antibiotic use in my dairy cattle/calves leads to bacterial infections in humans that are more difficult to treat. (3) | o | o | o | o | o | o |
| I would be willing to treat my dairy cattle/calves with antibiotic alternatives if they were equally effective. (4) | o | o | o | o | o | o |

Q6.24 What knowledge gaps need to be filled to make a raised without antibiotics production system more successful/sustainable and to have fewer unintended consequences?

________________________________________________________________

________________________________________________________________

________________________________________________________________

________________________________________________________________

________________________________________________________________

**End of Block: Beef**
